# Aging and false memories: Comparing effects of item-relatedness and list position

**DOI:** 10.1101/2025.07.28.666819

**Authors:** L. Gurguryan, E.A. Kensinger, P.A. Reuter-Lorenz, H.R. Dimsdale-Zucker

## Abstract

Evidence from classic serial position curves suggests that semantic and phonological false memories traditionally arise from presumed long and short-term memory stores, respectively. However, research challenges this distinction, finding semantic and phonological false memories across memory stores and list positions. The present study tested whether semantic and phonological false memories reflect distinct or shared mechanisms and whether these patterns differ with age. Younger (YAs) and older (OAs) adults completed a modified Deese-Roediger-McDermott paradigm in both recall (Experiments 1 and 2) and item recognition (Experiment 3), manipulating semantic and phonological similarity across list position. During recall, although OAs recalled fewer items than YAs, both groups generated more phonological than semantic false memories. During recognition, both groups had higher hit rates for phonological than semantic items and lower false alarm rates for the phonological compared to the semantic condition. Across all experiments, both groups made semantic and phonological false memories from all sublist positions, challenging notions of strictly separated memory stores. Importantly, irrespective of test format, there were no age-related differences in false memory patterns. Together, these findings challenge traditional views of memory organization and suggest that shared cognitive processes across age group and list composition underlie semantic and phonological false memories.

## Introduction

Although memory is often thought of as a factual recapitulation of our experiences, extensive laboratory work has shown that, instead, memory is prone to distortions and confabulations (Loftus, 1997). These inaccuracies, which can be measured in the lab as false memories, underscore the dynamic and reconstructive nature of memory (Bartlett, 1993; Schacter, 2012) and have been extensively studied using the Deese-Roediger-McDermott paradigm (DRM; Deese, 1959; Roediger & McDermott, 1995); for a review, see Gallo, 2010). In the canonical DRM paradigm, participants study 12-to-15 words semantically related to a non-presented theme (e.g., words like ‘*bed*,’ ‘*rest*,’ and ‘*pillow*’ are associated with the theme ‘*sleep*’), a list length that exceeds the canonical capacity of short-term memory (Miller, 1956). After encoding, participants freely recall the studied items from memory or complete a recognition memory task, often erroneously recalling or endorsing non-presented words that are semantically related to the theme word (e.g., ‘*sleep*’ in the example above; Deese, 1959; McDermott, 1996; McDermott & Roediger, 1998; Robinson & Roediger, 1997; Roediger & McDermott, 1995; Seamon et al., 1998; for a comprehensive review, refer to Gallo, 2013).

These memory errors, or false memories, are thought to arise from the automatic activation of related semantic lexical networks during encoding, whereby studied words activate related but unstudied concepts and words from long-term memory stores. In turn, these activated items interfere during remembering, producing false memories (Roediger et al., 2001). Given that this phenomenon depends on spreading activation within the long-term memory system, false memories have been traditionally been viewed as a “sin” of long-term memory (Gallo, 2013; Schacter, 2022)

However, accumulating evidence challenges the view that false memories are restricted to long-term memory stores. Evidence demonstrates that false memories can arise after relatively short retention delays and short lists of memoranda, which fall within the capacity of short-term memory stores. For instance, Atkins and Reuter-Lorenz (2008) found that participants falsely recognized semantically related lures after studying 4-item lists and being tested within seconds of encoding. Flegal and colleagues (2010) extended this work by testing false memories under both short- and long-term retrieval conditions and found that false memories occurred at similar rates in both conditions. Furthermore, participants rated true and false memories as subjectively similar in either the short-term or long-term version of the task. Festini & Reuter-Lorenz (2013) demonstrated that false memories can occur even with short 3- or 4-item lists and immediate testing with no retention delay, suggesting that short-term memory stores can support the generation of false memories (see also Abadie & Camos, 2019; Atkins & Reuter-Lorenz, 2008; Coane et al., 2007; Dimsdale-Zucker et al., 2018; Flegal et al., 2010). Together, these results suggest that common cognitive mechanisms may be at play during the generation of false memories regardless of whether retrieval is from short- or long-term memory stores.

Much of this research, however, involves different experimental conditions to elicit false memories in the short- versus long-term. To address this limitation, Dimsdale-Zucker and colleagues (2018) used a modified serial position curve paradigm to investigate short- and long-term false memories within a single task and recall window (see also Flegal et al. 2010 for use of a within-subjects design with identical study conditions across immediate and delayed tests). In this paradigm, participants studied 12-item lists with three distinct nested 4-item “sublists”. Each sublist consisted of four words that were semantically related to a unique theme (e.g., *‘hive’, ‘bumble,’ ‘sting,’ and ‘buzz’* are all semantically related to the theme *‘bee’*). Critically, in line with traditional views of serial position effects (Glanzer & Cunitz, 1966; Murdock, 1962), items from the early sublists are assumed to rely more heavily on long-term memory stores, while items from the most recent sublist are assumed to rely more on short-term memory stores (although see Waugh & Norman, 1965 for how contributions from long-term stores may contribute to recall in any list position). This design allowed false memories from both memory stores to be assessed concurrently within the same task and individual. Using this modified serial position paradigm, Dimsdale-Zucker and colleagues (2018) showed that false memories occurred across all sublists, although they were less frequent for the most recent sublist, especially when the retention interval was unfilled. These findings suggest that similar processes give rise to memory distortions regardless of list position and putative long- or short-term memory store.

Fuzzy Trace Theory (FTT; Brainerd & Reyna, 2002; Reyna & Brainerd, 1995) provides a framework for understanding how these memory errors can arise, irrespective of the underlying putative memory store. According to FTT, memoranda are simultaneously stored as both a verbatim and a gist representation. The verbatim representation encodes specific surface-level details (e.g., phonological or acoustic information) whereas the gist representation captures more general meaning and semantics. Together these representations support generally accurate memory. However, as verbatim traces tend to decay more rapidly, individuals increasingly rely on gist representations, which, while more durable, are also more prone to errors, and the likelihood of false memories.

This leads to items recalled from early portions of a list being preferentially supported by gist traces that are more resilient than fragile sensory coding that fades more rapidly (Brainerd & Reyna, 2002; Reyna & Brainerd, 1995). As a result, semantic, as compared to sound-based or phonological, false memories are more likely from earlier serial positions. Conversely, for later list positions, surface-level, sensory representations are presumed to be more prominent (Baddeley & Hitch, 1974), increasing the likelihood of sound-based or phonological confusions or false memories (Brainerd & Reyna, 2002; Reyna & Brainerd, 1995). Consistent with this, studies demonstrate that items in the primacy position are predominantly susceptible to semantic errors (Craik & Levy, 1970), while items in the recency position are selectively susceptible to acoustic or phonological errors (Craik, 1968).

Phonological false memories are also more common when there is a short rather than a long delay between encoding and retrieval (Lim & Goh, 2019; McBride et al., 2019 but see Atkins & Reuter-Lorenz, 2008), likely reflecting retrieval from short-term memory stores (Coane et al., 2024). However, in contrast with these predictions, Dimsdale-Zucker and colleagues (2018) found phonological false memories across all list positions. Albeit, these occurrences were rare given that study lists were constructed based on semantic similarity and were therefore intentionally biased to elicit semantic false memories. Complementing this, when lists are constructed based on phonological, as compared to semantic similarity, false memories can occur in the phonological domain at similar rates as seen in semantically related lists (e.g., recalling ‘*mat’* instead of ‘*cat’* or ‘*bat’*; (McBride et al., 2019; Sommers & Lewis, 1999; Watson et al., 2003).

Healthy aging offers a valuable lens through which to examine how differences in representational format influence false memory formation. Within FTT (Brainerd & Reyna, 2002; Reyna & Brainerd, 1995), age-related memory changes are often characterized as diminished access to verbatim traces alongside a greater reliance on gist-based processing. Consequently, as verbatim representations support the retrieval of item-specific details that can be used to reject related lures, their decline may increase older adults’ susceptibility to false memories.

Consistent with this, research shows that older adults generally recall less than their younger counterparts (Brickman & Stern, 2009; Craik & Salthouse, 2011; Rönnlund et al., 2005) and their recollections tend to be more reliant on gist-based traces rather than perceptual, verbatim traces (Abadie & Camos, 2019; Brainerd et al., 2008; Grilli & Sheldon, 2022; Schacter et al., 1997). Furthermore, on DRM tasks, younger adults generally outperform older adults, with older adults correctly reporting fewer studied items and also generating more semantic false memories as compared to younger adults (Balota et al., 1999; Budson et al., 2003; Norman & Schacter, 1997; Tun et al., 1998). Even when cues are provided that inform individuals about the nature of the DRM paradigm, false memories in older adults persist (McCabe & Smith, 2002). These findings suggest that aging is associated with an increased reliance on gist-based representations, which may selectively elevate semantic false memories. However, relatively little is known about how aging affects phonological false memories, particularly in paradigms that directly contrast semantic and phonological similarity. Recent work further suggests that different types of false memories may rely on distinct underlying mechanisms, as susceptibility to false memories across paradigms is often weakly related within individuals (Devitt & Foster, 2025). This highlights the importance of examining multiple forms of false memory within the same task and same participant.

An additional potential source of elevated false memory in older adults could arise from reduced availability of item-specific perceptual representations that would otherwise support discrimination and rejection of lures (Dennis et al., 2008; Naveh-Benjamin, 2000). Aging is associated with declines in sensory processing, including reductions in vision and hearing, with research showing that the degree of these sensory declines is linked to rates of cognitive decline (Roberts & Allen, 2016; for review, see Monge & Madden, 2016). The Information Degradation Hypothesis posits that sensory degradation leads to errors during perceptual processing, ultimately resulting in poor performance on higher-order cognitive tasks that may not necessarily be perceptual in nature (Schneider & Pichora-Fuller, 2000). Following this, one possibility is that declines in sensory processing contribute to the higher rates of false memories observed in older adults. Consistent with this, research indicates that older adults experience declines in memory precision, particularly the ability to discriminate between perceptually similar stimuli (Ferguson et al., 1992; Korkki et al., 2020; Stark et al., 2013; Toner et al., 2009; Yassa et al., 2011).

As such, under this alternative, but not mutually exclusive, account, weaker encoding or reduced memory precision would impair the ability to discriminate between studied items and similar lures, increasing susceptibility to false memories more broadly. Such reductions may stem from weaker initial encoding (Glisky et al., 2001) or difficulties in binding contextual details (Naveh-Benjamin, 2000; Old & Naveh-Benjamin, 2008).

Together, these accounts generate distinct predictions regarding the nature of age-related differences in false memory: If older adults primarily rely on gist-based processing, age differences should be most pronounced for semantically related lures. In contrast, if reduced access to item-specific detail is the primary driver, older adults should show elevated semantic and phonological false memories. Importantly, these effects may further interact with serial position, such that age differences may vary depending on whether retrieval is supported predominantly by stronger or weaker access to verbatim information.

To provide a rigorous test of these predictions, we adapted the paradigm used by Dimsdale-Zucker and colleagues (2018) and constructed lists where words either shared semantic or phonological associations with a non-presented theme (see **Appendix A**). Specifically, for half of the trials, the four words in each sublist were *semantically* related to a non-presented theme word (e.g., ‘*bed’*, ‘*nap’*, ‘*awake’*, ‘*doze’* for the theme word ‘*sleep’*). For the other half of the trials, the four words in each sublist were *phonologically* related to a theme word (e.g., ‘*leap*, ‘*steep*, ‘*bleep*, ‘*heap* for the theme word ‘*sleep’*). We operationalized false memories as the recollection of words corresponding to the preselected theme word, or words similar in meaning or sound but not included in the original study list, providing a parsimonious account of false-memory formation consistent with prior work (Atkins & Reuter-Lorenz, 2008; Dimsdale-Zucker et al., 2019; MacDuffie et al., 2012). This design allowed us to examine false memory generation across serial positions both within individuals (i.e., examining differences in semantic vs. phonological false memories across list positions) and between age groups (i.e., comparing performance between younger and older adults). In Experiment 1, younger adults (YAs) and older adults (OAs) studied and later recalled a series of word lists through simultaneous visual and auditory presentation. In Experiment 2, we replicated this in an OA-only sample that studied word lists via visual-only presentation. Experiment 3 extended this approach by examining memory in both OAs and YAs for the same types of studied items using an old/new recognition task, allowing us to determine whether effects of semantic and phonological similarity observed during recall also emerged when memory was assessed through recognition.

Based on extant theories, we predicted two possible patterns of results: First, based on accounts that posit memory for early and late study positions is supported by different underlying memory stores (Barnhardt et al., 2006; Brainerd & Reyna, 2002; Deese & Kaufman, 1957; Glanzer & Cunitz, 1966; Murdock, 1962; Reyna & Brainerd, 1995), we anticipated that semantic false memories would be more frequent for items presented in the early list positions—reflecting strong gist traces but relatively weaker verbatim traces. In contrast, we hypothesized that phonological false memories would be more likely for items presented at the end of the list where both gist and verbatim traces are presumed to be strong. Within this framework, we predicted that older adults, who exhibit reduced memory precision and impoverish access to item-specific detail, would generate more semantic rather than phonological false memories, even for items presented at later list positions.

Alternatively, if representations for early and late sublists instead rely on shared underlying memory stores and processes (Jonides et al., 2008; Nairne, 2002; Öztekin et al., 2010), we instead expected to observe similar rates of false memories across list positions and age groups. Specifically, we hypothesized that even if there were differences in the number of false memories from different sublist positions (e.g., more veridical and therefore fewer false memories from the most recent positions), that there would be no observable differences in the type of false memories—semantic or phonological—from these different sublists for both YAs and OAs. Altogether, this study aims to contribute to the ongoing discussion about the dual- versus single-store nature of underlying memory stores that can support both successful and erroneous memory performance. By examining both recall and recognition, the present experiments further allow us to assess whether effects of semantic and phonological similarity extend across different measures of memory performance, thereby providing new insights into the theoretical understanding of false memories and the structure of memory itself.

## Methods

### Experiment 1

#### Participants

Native English speakers, free from any reported neurological or psychological disorders, and naïve to the stimulus set were recruited for this study. Forty-eight Boston College undergraduate students (30 = female) participated either for course credit (0.5 credits/half hour) or cash ($5/half hour). All participants completed a battery of neuropsychological assessments that included the Mini-Mental State Examination (MMSE; Folstein et al., 1975), the Shipley Institute of Living Scale (SILS; Shipley, 1986), and the FAS (total, perseverations, and intrusions) to evaluate cognitive function, memory, and verbal fluency. The Digit Symbol Substitution Test, Digits Forward, and Digits Backward tasks were used to assess processing speed and working memory. The Wisconsin Card Sorting Test (WCST) was administered to measure executive function and cognitive flexibility. The Beck Anxiety Inventory (BAI) and Beck Depression Inventory (BDI; Beck et al., 1996) were used to evaluate levels of anxiety and depression. Given that depression is associated with deficit in episodic memory (for review, see James et al., 2021), six participants were excluded for scores greater than 10 on the BDI (per scores <10 being indicative of mild to moderate depression; Beck et al., 1988), leaving a remaining sample of 42 participants (25 = female; Mean_age_= 19.34 years, SD_age_= 0.99 years; see **Table 1** for neuropsychological assessment data).

**Table 1.**
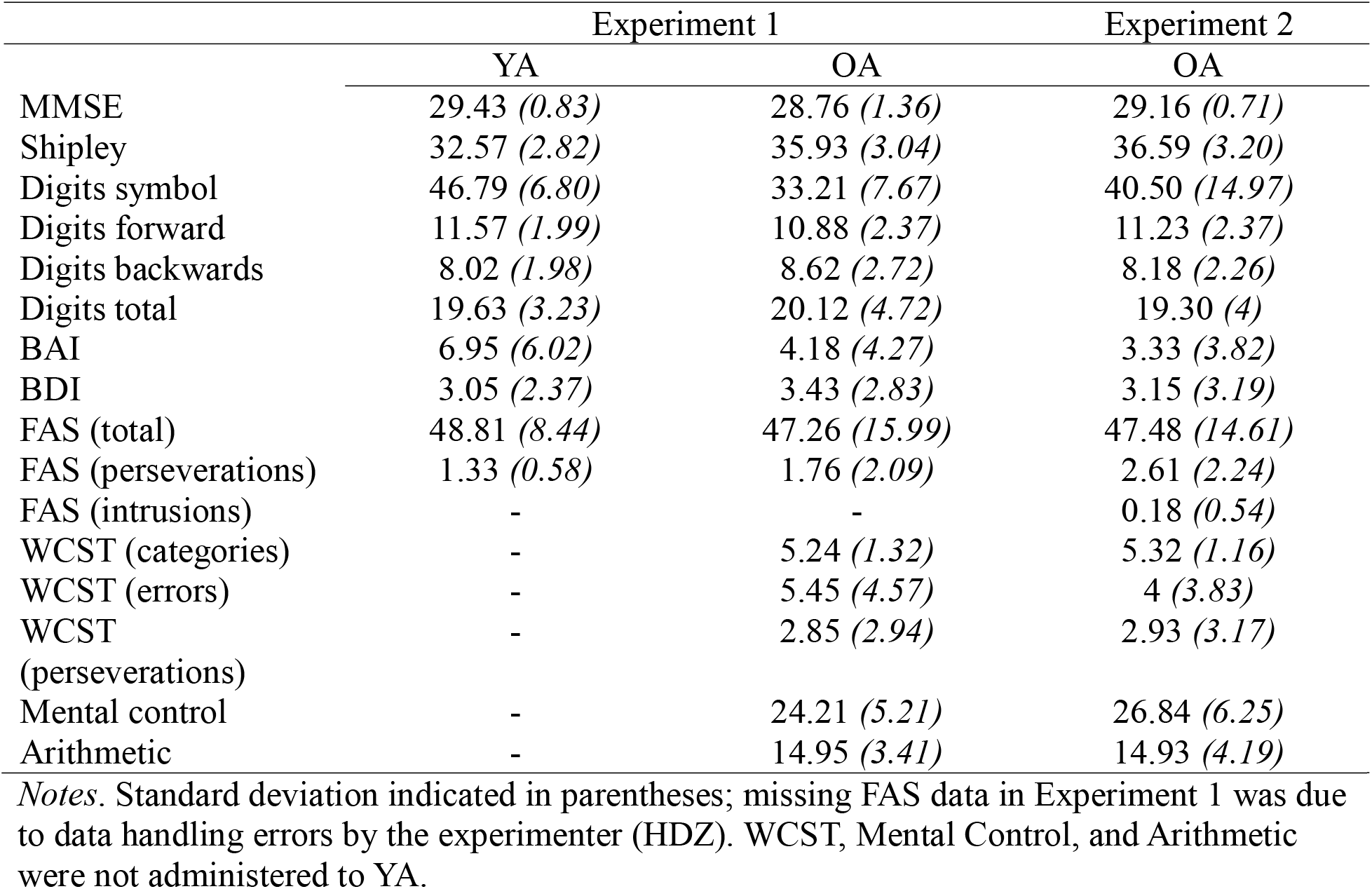
Neuropsychological assessment data.

|  | Experiment 1 |  | Experiment 2 |
| --- | --- | --- | --- |
|  | YA | OA | OA |
| MMSE | 29.43 (0.83) | 28.76 (1.36) | 29.16 (0.71) |
| Shipley | 32.57 (2.82) | 35.93 (3.04) | 36.59 (3.20) |
| Digits symbol | 46.79 (6.80) | 33.21 (7.67) | 40.50 (14.97) |
| Digits forward | 11.57 (1.99) | 10.88 (2.37) | 11.23 (2.37) |
| Digits backwards | 8.02 (1.98) | 8.62 (2.72) | 8.18 (2.26) |
| Digits total | 19.63 (3.23) | 20.12 (4.72) | 19.30 (4) |
| BAI | 6.95 (6.02) | 4.18 (4.27) | 3.33 (3.82) |
| BDI | 3.05 (2.37) | 3.43 (2.83) | 3.15 (3.19) |
| FAS (total) | 48.81 (8.44) | 47.26 (15.99) | 47.48 (14.61) |
| FAS (perseverations) | 1.33 (0.58) | 1.76 (2.09) | 2.61 (2.24) |
| FAS (intrusions) | - | - | 0.18 (0.54) |
| WCST (categories) | - | 5.24 (1.32) | 5.32 (1.16) |
| WCST (errors) | - | 5.45 (4.57) | 4 (3.83) |
| WCST (perseverations) | - | 2.85 (2.94) | 2.93 (3.17) |
| Mental control | - | 24.21 (5.21) | 26.84 (6.25) |
| Arithmetic | - | 14.95 (3.41) | 14.93 (4.19) |

*Notes.* Standard deviation indicated in parentheses; missing FAS data in Experiment 1 was due to data handling errors by the experimenter (HDZ). WCST, Mental Control, and Arithmetic were not administered to YA.

An additional sample of 46 healthy, community dwelling seniors (aged 59 and up) were recruited from the Greater Boston area and compensated with cash ($5/half hour) for their participation. One participant was excluded for scores greater than 10 on the BDI (Beck et al., 1996). An additional 3 participants withdrew themselves following the practice citing task difficulty and personal concerns about memory abilities precluding completion of the task. Results presented below are from the remaining 42 older participants (27 = female; Mean_age_= 76.31 years, SD_age_= 6.70 years; see **Table 1** for neuropsychological assessment data).

While an a priori sample size calculation was not conducted, the sample size of 42 participants per age group was determined based on precedent from prior work from our group (Dimsdale-Zucker et al., 2018), which successfully detected relevant effects with a comparable design. We additionally conducted a post hoc sensitivity analysis to estimate the smallest effect size detectable with our sample (N = 42). Although such analyses do not substitute for an a priori power analysis, they can help contextualize the study’s sensitivity. Using G*Power (Faul et al., 2007), we determined that with 42 subjects per group, six conditions (2x3), α = .05, and power = .80, the smallest detectable effect size was Cohen’s f = 0.228 (η² = .049), corresponding to a small-to-medium effect. This estimate, based on a conservative one-way ANOVA approximation rather than the exact repeated-measures design used here, suggests that the study was sufficiently sensitive to detect small-to-medium effects.

#### Materials

One 12-item list was presented on each of 46 trials (see **Appendix A**). Words were printed on a computer screen while a computerized voice simultaneously read the words aloud. Each list was composed of three sublists where each sublist contained four words that were all related to a common non-presented theme word (see **Appendix B**). On half of the trials, associates were related to the theme semantically (see **Figure 1B**) and for the other half of trials phonologically (see **Figure 1B**). Order of semantic and phonological trials was randomized for each participant. Semantic versus phonological relationship of list items was counterbalanced across participants such that each triplet was only presented once in either the semantic or phonological condition for each participant. Additionally, the order of sublists in each list were counterbalanced so that each theme occurred in each presentation position (primacy, middle, recency) equally often across participants.

**Figure 1.**
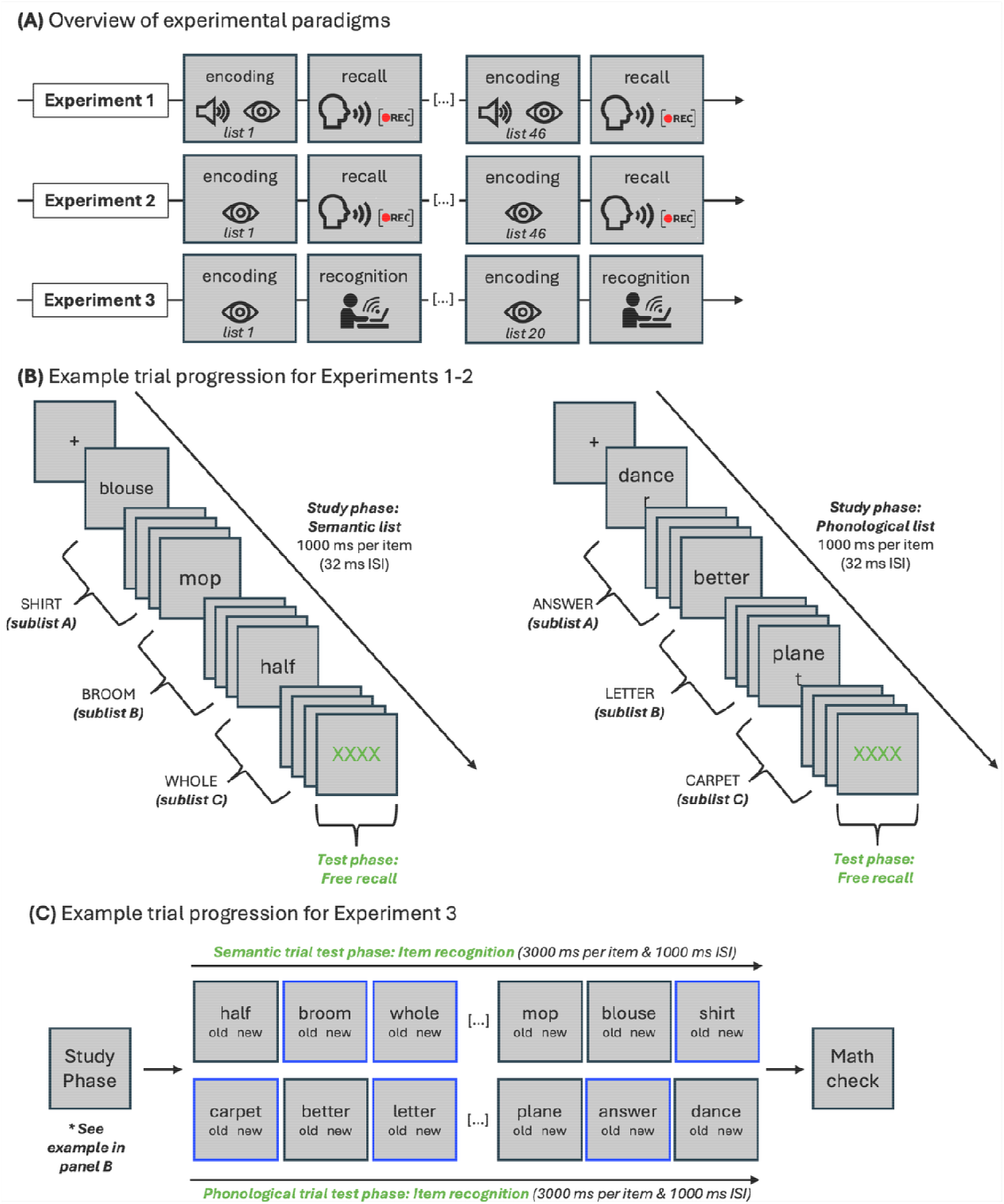
**(A)** Overview of the experimental paradigms for Experiments 1 and 2, which were conducted in person, and Experiment 3, which was conducted online. **(B)** Example trial progression of the study and test (i.e., free recall task) phases for Experiment 1 and 2. **(C)** Example trial progression of the study and test phase (i.e., item recognition task) for Experiment 3; blue outlines denote new items during the test phase and are for illustration purposes only.

Sublists represent a subset of 128 lists taken from previously published work on semantic false memories in STM (Flegal et al., 2010). These lists were supplemented with additional themes taken from neutral associates published in Kensinger and Corkin (2003) and associates were selected from the University of South Florida (USF) Free Association Norms (Nelson et al., 2004) following similar procedures as previous work (Flegal et al., 2010) to yield a total of 138 themes that were organized into 46 possible triplets. Audio files for each word were culled from the Linguistic Data Consortium at UPenn (http://www.ldc.upenn.edu/). Additional stimuli not available from this dataset were constructed in Lips (http://mac-free.com/download/Lips.html) using the female “Victoria” voice and further edited in Audacity (http://audacity.sourceforge.net/).

#### Procedure

Participants gave written consent in accordance with the Boston College IRB. The experimenter explained the memory task and then participants completed five practice trials before proceeding through the remaining 46 task trials. Experimenters remained in the room and transcribed audio responses on laptop computers. Audio responses were additionally recorded using an Olympus WS-210s voice recorder to allow for later analysis. Following the memory task, participants were given feedback about the task, completed neuropsychological tasks, and were debriefed as to the purpose of the study.

Words were presented serially at a rate of 1000 ms per word (after Bartha et al., 1998) with 32 ms between words. Timing and presentation were controlled using ePrime 2.0 software (Psychology Software Tools, Pittsburgh, PA). Stimuli appeared in black Courier New font on a silver background. Presentation of the 12 words was followed by a row of five green “X”s that served as the recall cue and remained the screen for the duration of the free recall. Participants were required to take a minimum of 30 seconds to freely recall the words aloud. Participants pressed the space bar to advance to the next set of words which began 1000 ms later.

#### Data analysis

In addition to the experimenter’s in-room transcriptions, a second experimenter transcribed each audio file. A third coder resolved discrepancies between these transcriptions. Incorrect responses were classified as (1) *semantic:* associates listed in the USAF norms as related to the theme word or judged as related in meaning to the memory set by a trained coder^1^, and (2) *phonological:* related in sound to the theme word. All other error types—including within-trial repeats, between-trial repeats, unrelated responses, and non-words or mispronunciations—were collapsed into a miscellaneous error category (see Atkins & Reuter-Lorenz, 2008 for similar scoring procedures). As no a priori hypotheses concerned the relation of these miscellaneous errors to sublist or condition, this category was excluded from the primary analyses. Throughout the manuscript, degrees of freedom of the ANOVAs were corrected using the Geisser-Greenhouse method to adjust for violations of sphericity, resulting in non-integer values for the degrees of freedom and Tukey’s HSD (Honestly Significant Difference) was used to ensure accurate pairwise comparisons.

### Results

#### Total number of recalled words (combined correct and false)

A repeated measures ANOVA with condition (semantic, phonological) and sublist (A, B, C) entered as a within-subjects factor and age group (YA, OA) specified as a between-subjects factor indicated a significant main effect of age group (*F*(1, 82) = 34.66, *p* < 0.001, ^2^ = 0.30) such that YAs recalled overall a greater number of words as compared to OAs (see **Table 2** for numeric values). There was a significant main effect of condition (*F*(1, 82) = 134.82, *p* < 0.001, = 0.62) showing that participants recalled a greater number of words in the semantic relative to the phonological condition. The main effect of sublist was also significant (*F*(1.63, 133.56) = 297.56, *p* < 0.001,*η*²_p_= 0.79). Participants recalled a greater number of words in sublist A compared to sublist B (*t*(82) = 2.49, *p* = 0.04); however, they recalled significantly fewer words in sublist A (*t*(82) = −16.55, *p* < 0.001) and in sublist B (*t*(82) = −23.83, *p* < 0.001) compared to sublist C. This analysis also revealed a significant interaction effect between age group and condition (*F*(1, 82) = 35.97, *p* < 0.001,*η*²_p_ = 0.31). Both YA (*t*(82) = −12.45, *p* < 0.001) and OA (*t*(82) = −3.97, *p* = 0.001) participants reported a greater number of words in semantic compared to phonological trials, though the magnitude of this effect was stronger in YAs. None of the other effects—specifically, the interactions between age group and sublist, condition and sublist, nor the three-way interaction between age group, condition, and sublist—reached statistical significance (all *p*-values > 0.05). See **Figure C1A** in **Appendix C** for an illustration of the data.

**Table 2.**
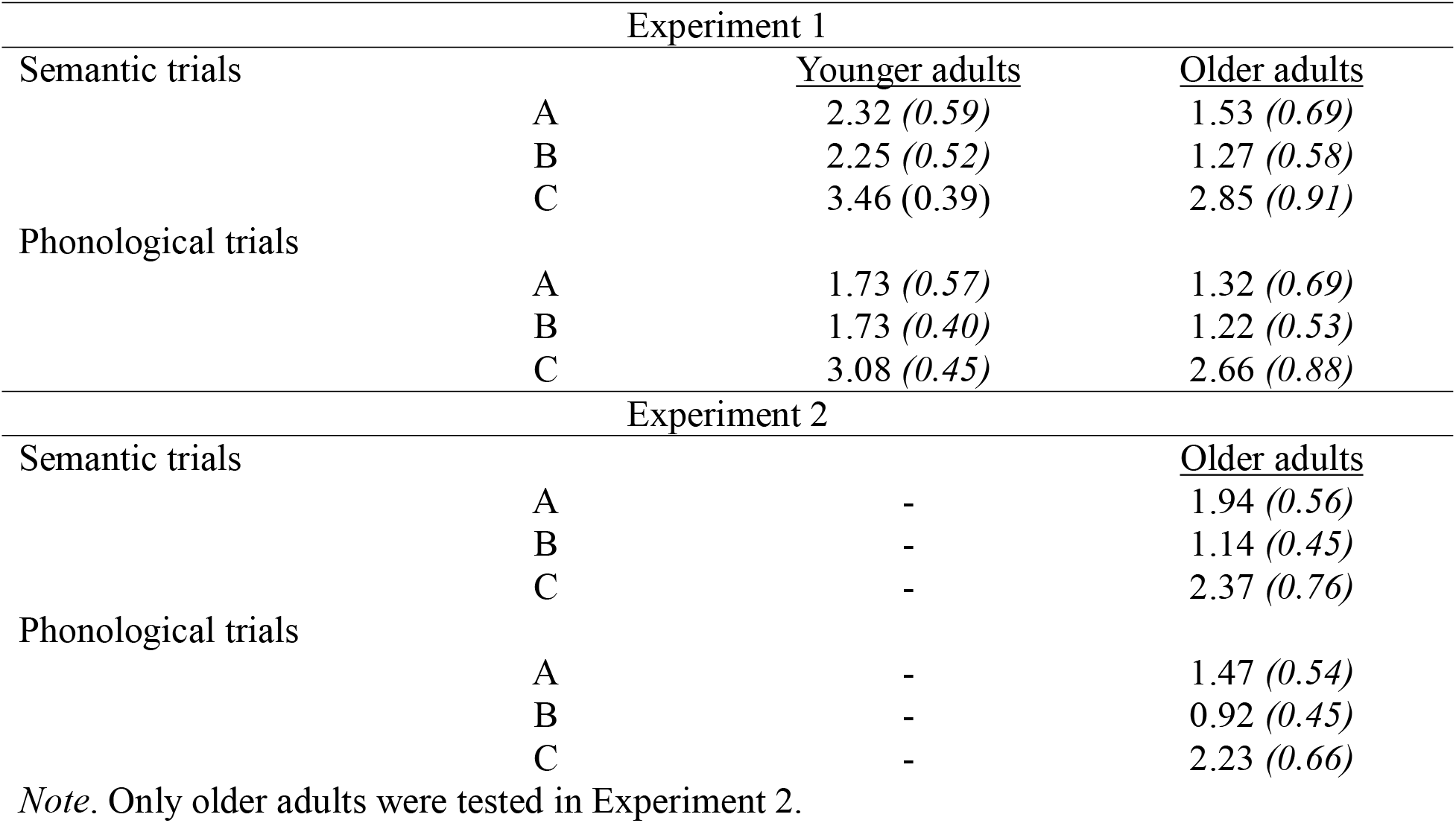
Total number of recalled words (correct and false combined; means with standard deviation in parentheses) split by age group, list type, sublist, and experiment.

#### Correct responses (output accuracy)

To account for the overall differences in the total number of items recalled between YAs & OAs, we re-scored the number of correct responses as a proportion of total recalled items. This approach—commonly used when analyzing data across different age groups (e.g., Khanna & Cortese, 2009; McGuire et al., 2015; Sugrue & Hayne, 2006)—enables us to remove age-related differences in overall recall rates. Of note, our calculation reflects the relative balance of correct versus false memories in each recall attempt (i.e., output accuracy) rather than relative to total items presented. This approach enables comparison of the distribution of correct versus false memory responses within participants’ output. For completeness, we report raw descriptive statistics of the number of correct and false memory responses without this scoring correction in **Table 3**. Additionally, we computed and report serial position curves (Murdock, 1962) to examine the probability of correct responses as a function of presentation position. These curves revealed a recency effect such that there was a greater probability of correct responses from the last list position relative to the first and middle lists (see **Figure 2**); this was replicated with statistical analyses (reported below).

**Figure 2.**
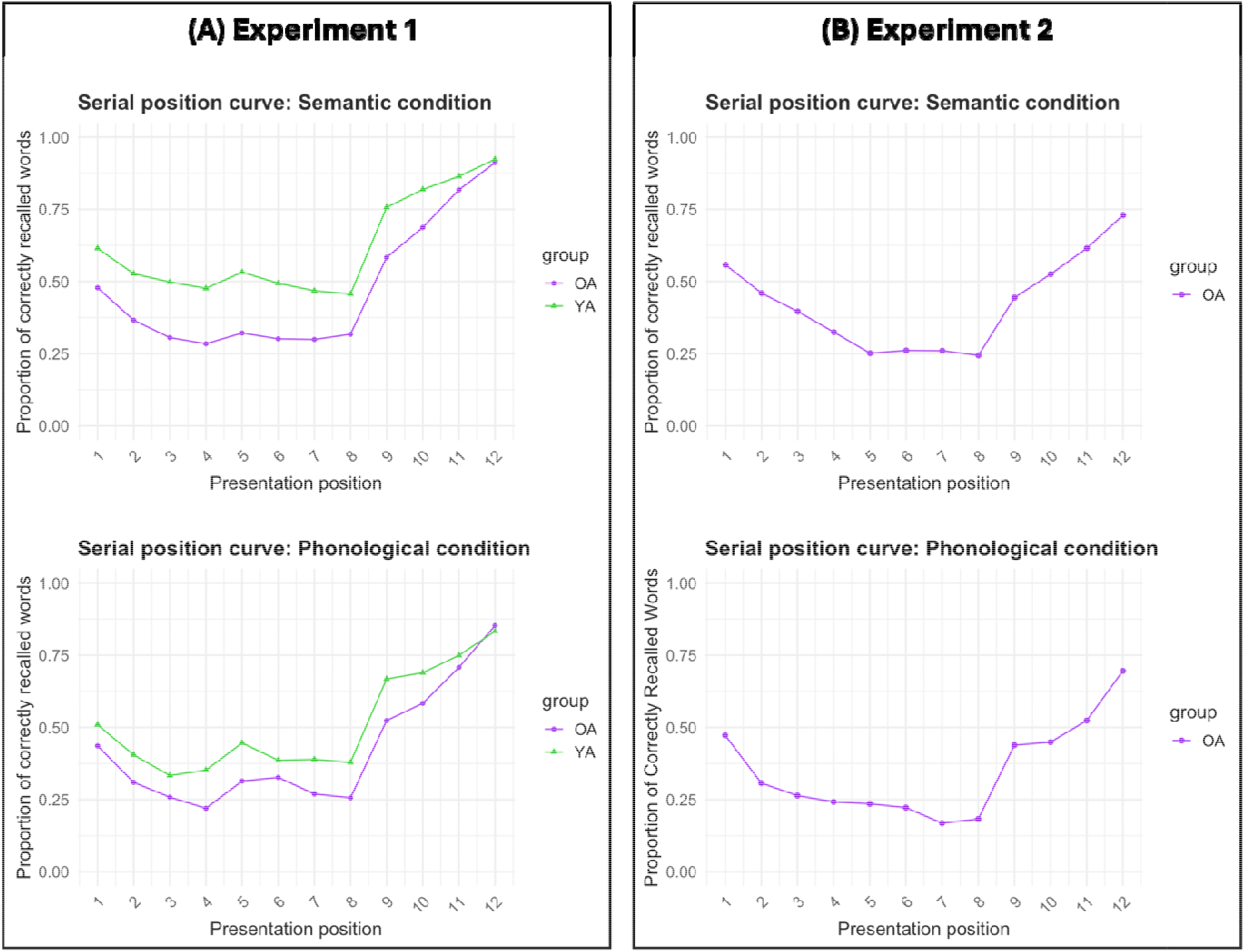
Serial position curves (correct responses) as a function of condition (semantic, phonological) for **(A)** Experiment 1 and **(B)** Experiment 2 (only older adults were tested in Experiment 2).

**Table 3.**
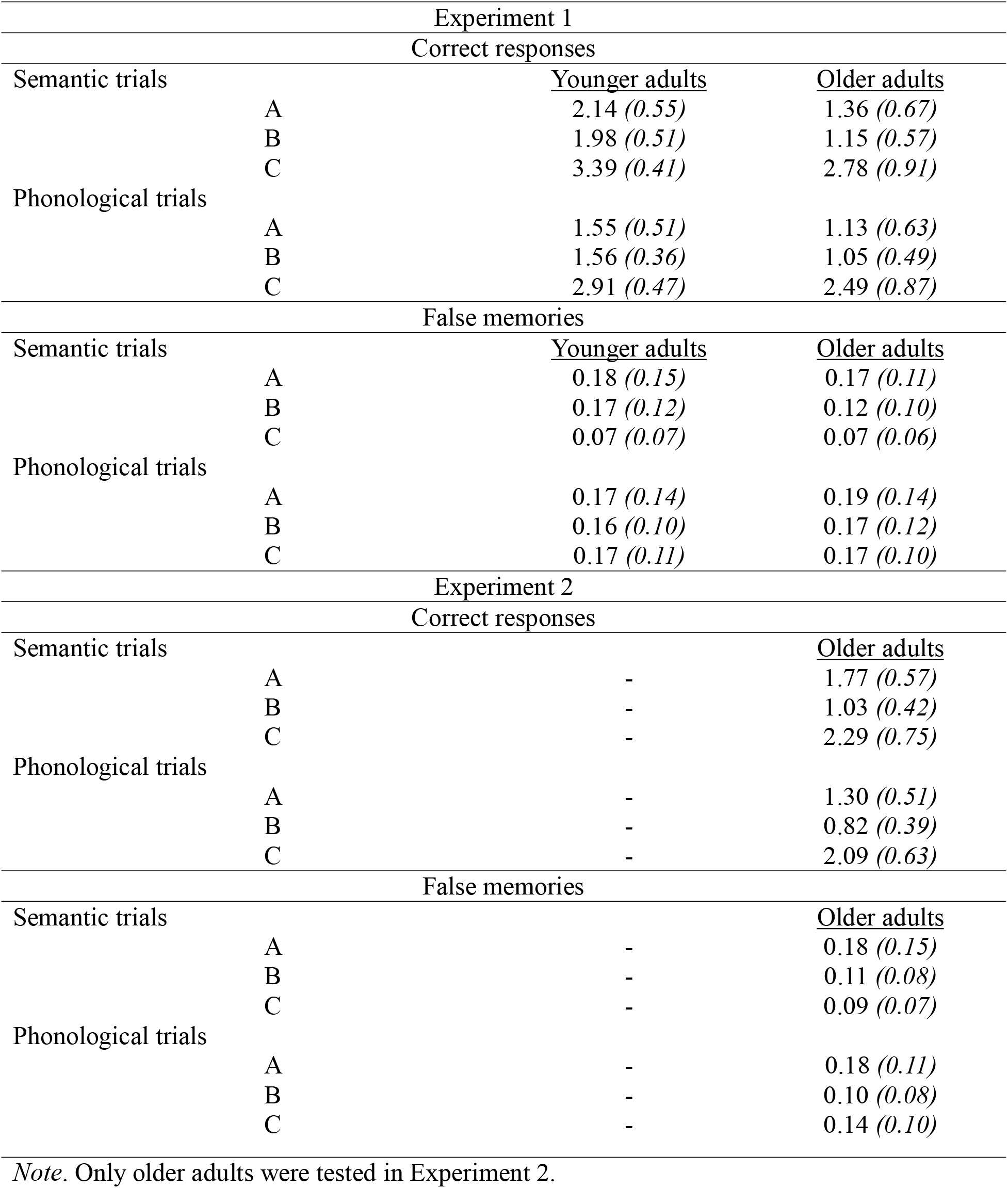
Raw means of correct words and false memories recalled (standard deviation in parentheses) split by age group, list type, sublist, and experiment.

A repeated measures ANOVA was conducted with condition (semantic, phonological) and sublist (A, B, C) entered as within-subjects factors and age group (YA, OA) as a between-subjects factor. There was a significant main effect of age group (*F*(1, 82)= 5.03, *p* = 0.028, ^2^ = 0.06) such that YA reported a greater proportion of correct words relative to OA (*t*(82) = −2.24, *p* = 0.03). The main effect of condition was also significant (*F*(1, 82) = 21.85, *p* < 0.001, ^2^ = 0.21) such that there was a greater proportion of correctly recalled words in the semantic relative to the phonological condition (*t*(82) = 4.67, *p* < 0.001). Lastly, there was a significant main effect of sublist (*F*(1.93, 158.26) = 69.62, *p*< 0.001, ^2^ = 0.46) indicating significant differences between sublists A and C (*t*(82) = −9.86, *p* < 0.001) and sublist B and C (*t*(82) = −11, *p* < 0.001) while no significant differences were found between sublists A and B (*t*(82) = −0.55, *p* = 0.85). That is, participants reported a greater proportion of correct words in sublist C relative to both sublists A and B. All other effects, notably including an interaction between condition, sublist, and age, did not reach the threshold for statical significance (all *p* > 0.05). See **Table 4** for descriptive statistics and **Figure C2A** in **Appendix C** for an illustration of the data. Thus, although we observed numerically greater memory for the earliest presented items in the list, we failed to find a statistically significant primacy effect.

**Table 4.**
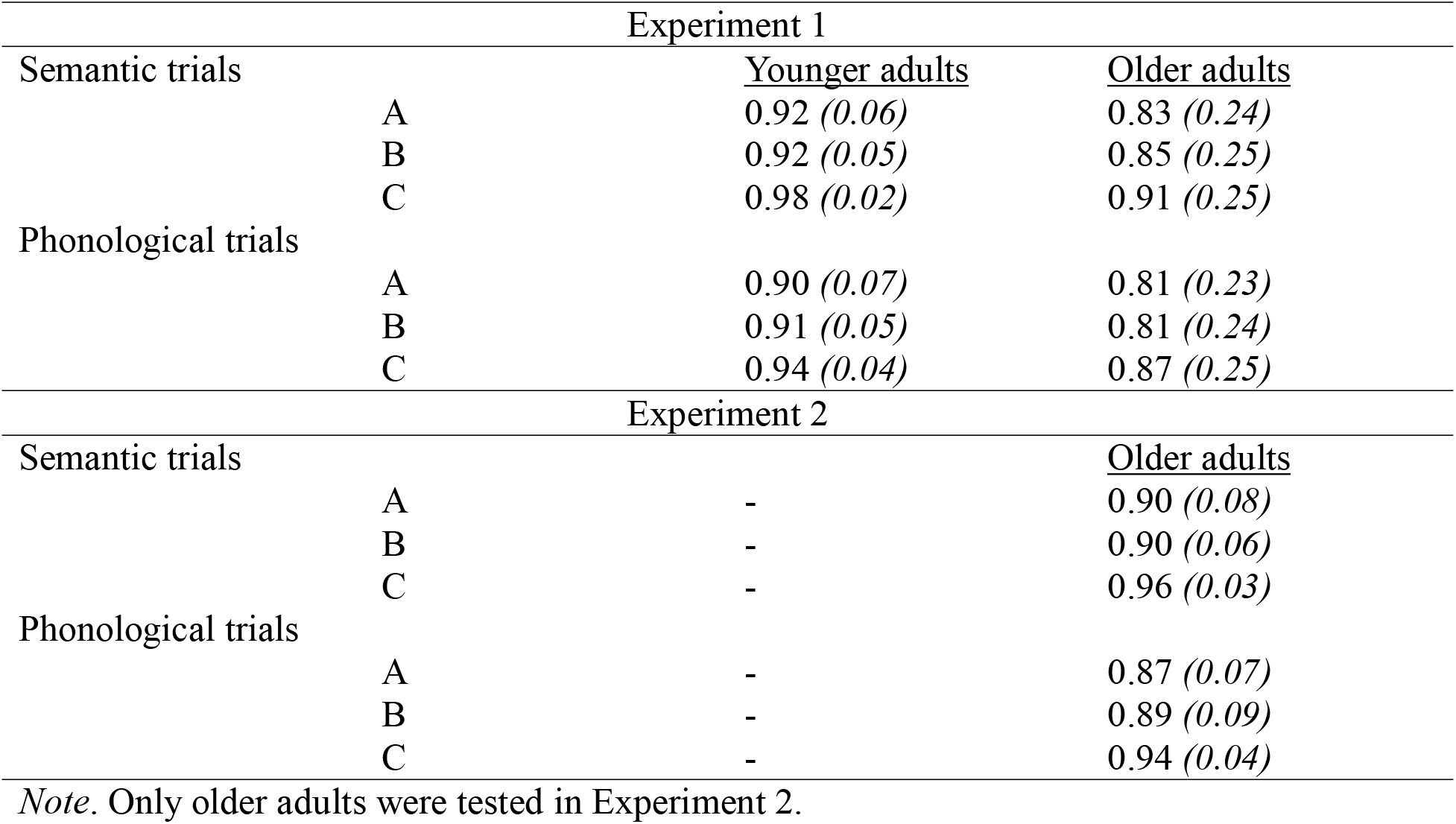
Proportion of correct words recalled (standard deviation in parentheses) split by age group, list type, sublist, and experiment.

#### False memories

A repeated measures ANOVA was conducted with condition (semantic, phonological) and sublist (A, B, C) entered as within-subjects factors and age group (YA, OA) entered as the between-subjects factor. This analysis revealed a significant main effect of group (*F*(1, 82) = 5.03, *p* = 0.03,*η*²_p_ = 0.06) such that, on average, OAs had an increased proportion of false memories as compared to YAs (*t*(82) = 2.24, *p* = 0.03). There was a significant main effect of condition (*F*(1, 82) = 21.85, *p* <0.001) indicating an overall greater proportion of false memories in the phonological relative to the semantic condition (*t*(82) = −4.67, *p* < 0.001). There was a significant main effect of sublist (*F*(1.93, 158.26) = 69.62, *p* < 0.001, ^2^ = 0.46), establishing significant differences between sublist A and sublist C (*t*(82) = 9.86, *p* <0.001), as well as between sublist B and sublist C (*t*(82) = 11, *p* < 0.001). Both sublists A and B showed higher proportions of false memories compared to sublist C. No other significant effects were found, notably including with age (all *p* > 0.05). See **Table 5** for descriptive statistics and **Figure 3A** for an illustration of the data.

**Figure 3.**
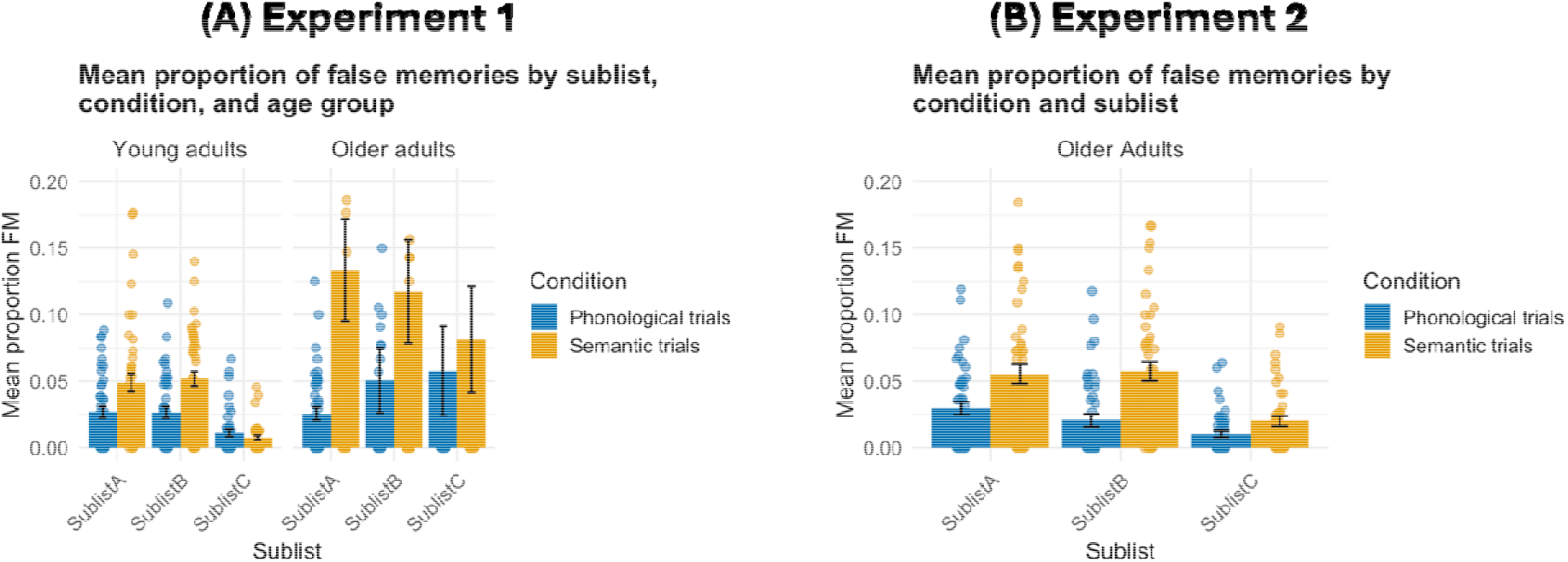
**(A)** Average proportions of false memories recalled as a function of age group (YA, OA), condition (semantic, phonological), and sublist (A, B, C). **(B)** Average proportions of false memories recalled as a function of condition (semantic, phonological), and sublist (A, B, C). In the semantic condition, false memories refer to semantic false memories, while in the phonological condition, they refer to phonological false memories. Only older adults were tested in Experiment 2. Points indicate individual participants and error bars indicate standard error of the mean.

**Table 5.**
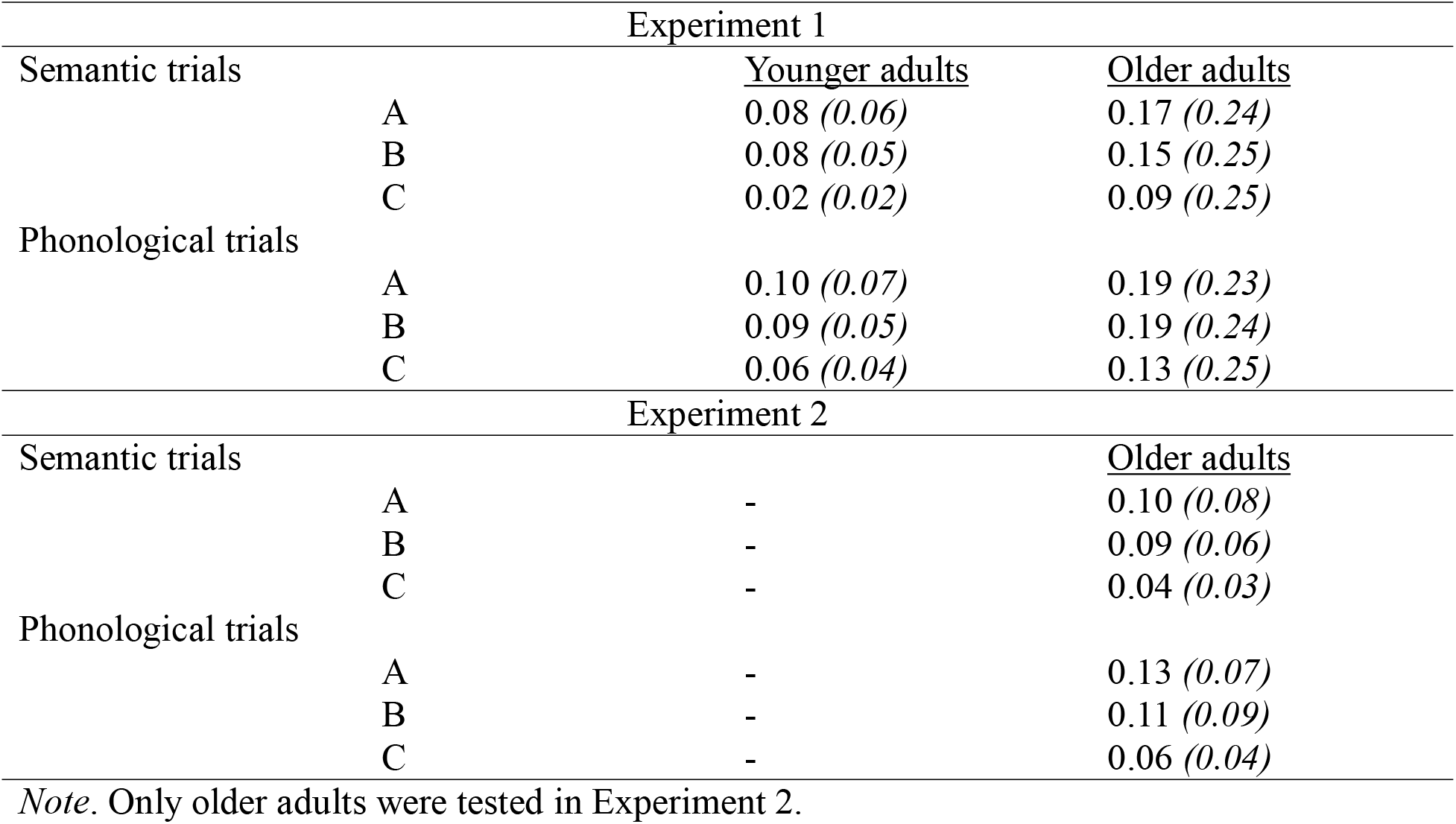
Proportion of false memories (standard deviation in parentheses) split by age group, list type, sublist, and experiment.

### Experiment 1: Summary

In Experiment 1, we examined performance on an adapted version of the DRM paradigm, focusing on the effects of list type (semantic vs. phonological) and sublist position (A, B, C) between age groups (YAs and OAs). Our findings revealed several key patterns across both age groups. All participants, irrespective of age, recalled a higher proportion of correct words in the semantic condition compared to the phonological condition, with this effect being more pronounced in YAs. A recency effect was also observed, as all participants recalled a higher proportion of correct words from sublist C, regardless of age or condition. Furthermore, all participants generated a lower proportion of false memories from sublist C, suggesting that verbatim traces may have been more robust and thus better able to counteract gist traces from this most recent sublist (Brainerd & Reyna, 2002; Reyna & Brainerd, 1995).

Interestingly, both groups exhibited similar patterns of false memory production. That is, despite OAs remembering fewer items overall relative to YAs, all participants generated a higher proportion of false memories in the phonological relative to the semantic condition and had similar distributions of false memories across the sublists (fewest false memories from the most recent C sublist than either sublist A or B). Importantly, these age-equivalent patterns emerged despite OAs generating more false memories overall, highlighting the robustness of these effects even in the context of significant main effects of age. This finding contradicts the prediction that OAs would be more prone to phonological errors, especially from the most recent C sublist, as compared to YAs, and suggests that false memory production is similar across age groups.

Our findings also challenge the view that the nature of false memories is strictly tied to a specific memory store, suggesting instead that the types of false memories generated are more closely related to the properties of the studied lists (for an account of how rehearsal may enable transfer between putative stores, see Waugh & Norman, 1965). To further investigate whether the surprising finding of comparable performance observed in OAs and YAs was influenced by the bimodal presentation of stimuli (i.e., both visual and auditory), we conducted Experiment 2, focusing solely on OAs in which we presented words visually only. Prior work has indicated that presenting information in multiple modalities may enhance task performance in older adults (e.g., Mahoney et al., 2011; for a review, see De Dieuleveult et al., 2017). As such, if the enhanced encoding from the bimodal presentation in Experiment 1 masked age-related differences, we expected that removing auditory cues would reduce memory performance in OAs.

## Experiment 2

### Methods

#### Participants

A sample of 47 healthy, community dwelling seniors (aged 59 and up) were recruited from the Greater Boston area and compensated at a rate with cash ($5/half hour) for their participation. Three participants were excluded: 2 participants were excluded for having an MMSE score less than 28 and 1 participant was excluded due to errors during data collection. Results presented below are from the remaining 44 older participants (26 = female; Mean_age_= 72.5 years, SD_age_= 7.11 years; see **Table 1** for neurological assessment data).

#### Materials

All materials were identical to those used in Experiment 1.

#### Procedures

All procedures were identical to those in Experiment 1, with the exception that stimuli were presented visually only, rather than presented both visually and auditorily.

#### Data analysis

All analyses were identical to those in Experiment 1.

### Results

#### Total number of recalled words (combined correct and false)

A repeated measures ANOVA with condition (semantic, phonological) and sublist (A, B, C) entered as within-subjects factors was conducted on the overall number of words recalled, regardless of correct or FM status. This analysis revealed, first, a significant main effect of condition (*F*(1, 43) = 105.33, *p* < 0.001,*η*²_p_= 0.7) showing that participants recalled on average a greater number of words in the semantic relative to the phonological condition (*t*(43) = 10.26, *p* < 0.001). Second, there was a significant main effect of sublist (*F*(1.37, 58.80) = 59.13, *p* < 0.001,*η*²_p_= 0.58). Pairwise comparisons indicated that participants recalled significantly more words in sublist A compared to sublist B (*t*(43) = 7.29, *p* < 0.001). Conversely, they recalled significantly fewer words in sublist A (*t*(43) = −3.90, *p* < 0.001) and in sublist B (t(43) = −13.12, p < 0.001) as compared to sublist C. Lastly, this analysis revealed a significant interaction effect between condition and sublist (*F*(1.96, 84.15) = 7.39, *p* < 0.001,*η*²_p_= 0.15). Pairwise comparisons indicated that within all three sublists (A: *t*(43) = 7.32, *p* < 0.001; B: t(43) 3.81, *p* < 0.001; C: *t*(43) = 2.68, *p* = 0.01), participants recalled more words in the semantic relative to the phonological condition. See **Table 2** for detailed descriptive statistics and **Figure C1B** in **Appendix C** for an illustration of the data.

#### Correct responses (output accuracy)

A repeated measures ANOVA with condition (semantic, phonological) and sublist (A, B, C) specified as within-subjects factors was conducted. While the interaction between the factors was not significant (*F*(1.66, 71.34) = 0.26, *p* = 0.73,*η*²_p_= 0.01), the analysis revealed a significant main effect of condition (*F*(1, 43) = 16.23, *p* < 0.001,*η*²_p_= 0.27). Participants recalled a greater proportion of correct words in the semantic relative to the phonological condition (*t*(43) = 4.03, *p* < 0.001). Additionally, the main effect of sublist (*F*(1, 98, 85.04) = 26.18, *p* <0.001, ^2^ = 0.38) was significant. While there was no significant difference in the proportion of correctly recalled words between sublist A and B (*t*(43) = −0.77, *p* = 0.73), participants recalled a significantly lower proportion of correct words in sublist A relative to sublist C (*t*(43) = −6.94, *p* < 0.001) as well as a significantly lower proportion of correct words in sublist B relative to sublist C (*t*(43) = −5.85, *p* < 0.001). See **Table 3** for descriptive statistics and **C2B** for an illustration of the data.

#### False memories

A repeated measures ANOVA with condition (semantic, phonological) and sublist (A, B, C) entered as within-subjects factors was conducted. This analysis revealed a main effect of condition (*F*(1, 43) = 16.23, *p* < 0.001, ^2^ = 0.27) such that participants reported a greater proportion of false memories in the phonological relative to the semantic condition (*t*(43) = −4.03, *p* < 0.05). The main effect of sublist was also significant (*F*(1. 98, 85.04) = 26.18, *p* < 0.001,*η*²_p_=0.38). Pairwise comparisons showed that while there was no significant difference in the proportion of false memories between sublists A and B (*t*(43) 0.77, *p* = 0.73), sublist C had a significantly lower proportion of false memories compared to both sublists A (*t*(43) = 6.94, *p* < .001) and B (*t*(43) = 5.85, *p* < .001). Lastly, the interaction between the factors was not significant (*F*(1.66, 71.34) = 0.26, *p* = 0.81, ^2^ =0.01). See **Table 5** for detailed descriptive statistics and **Figure 3B** for an illustration of the data.

## Experiment 2: Summary

In Experiment 2, we focused our investigation solely on OAs and replicated key findings from Experiment 1: First, OAs recalled a higher proportion of correct words in the semantic condition compared to the phonological condition, mirroring the pattern observed in Experiment 1. A recency effect was also evident, as in Experiment 1, with OAs recalling a higher proportion of correct words from sublist C. When analyzing false memory production, OAs generated a higher proportion of phonological false memories compared to semantic false memories. Additionally, a higher proportion of false memories were observed from sublist A relative to sublist C, complementing the finding that more items were correctly recalled from the end of the presented lists supporting the recency effect seen in correct recall, and from sublist B relative to sublist C; however, there was no interaction between the types of false memories generated and list position.

One notable pattern across the two experiments was that OAs exhibited numerically higher performance (i.e., increased accurate recall) in Experiment 2 than in Experiment 1. Although there were some demographic differences between the samples—e.g., OAs in Experiment 1 were somewhat older than those in Experiment 2—these are unlikely to fully account for the observed differences in recall performance. Instead, it appears that the bimodal (i.e., simultaneous auditory and visual) presentation format used in Experiment 1 may have been unexpectedly detrimental for OAs. While prior research suggests that multimodal presentation of information can enhance task performance in older adults (for a review, see De Dieuleveult et al., 2017), it is possible that the bimodal presentation in this case introduced additional processing demands that negatively affected OA performance on our task. Nevertheless, the patterns observed in Experiment 2 were largely consistent with those found in Experiment 1. This aligns with other research showing that presenting visual stimuli, with or without auditory input, results in similar effects on learning and memory recall in healthy older adults (Constantinidou & Baker, 2002).

However, Experiments 1 and 2 left open questions about how memory performance differs between YAs and OAs when additional retrieval support is provided. To address this question, we preregistered Experiment 3 (https://aspredicted.org/67k4i8.pdf; preregistration #304,917), which retained the unimodal encoding presentation used in Experiment 2 but provided additional retrieval support by changing the task from free recall to recognition memory. The recognition memory format also allowed us to directly examine study–test lag— defined as the number of intervening items between the original studied item and its corresponding lure at test— providing an opportunity to test whether the relationship between memory performance and lag differs across age groups and list type. Specifically, this analysis extended our previously reported sublist analysis by examining whether susceptibility to false memories varied as a function of the temporal distance between studied items and their corresponding lures and whether this relationship differed depending on whether the original item came from sublist A, B, or C.

### Methods

#### Participants

An a priori power analysis was conducted using the WebPower package in R for a 2 (age group: younger, older) x 2 (condition: semantic, phonological) mixed repeated-measures ANOVA. Assuming a medium effect size (*f* = 0.25), α = .05, and 80% power, the required total sample size was 64 participants per group. Participants were recruited through Prolific (https://www.prolific.com/) and were compensated at a rate of $6.00 per hour. To meet our a priori sample size, one hundred and thirty-four participants were tested, but 6 participants were excluded from analyses based on predetermined exclusion criteria described in the preregistration: 1 YA participant was excluded for scoring below 75% on the math attention check and 5 participants (3 YAs, 2 OAs) were excluded for missing more than 50% of responses throughout the task. The final sample includes 64 YAs (Mean_age_= 24.8 years, SD_age_= 4.84 years; 38 female, 22 male, 1 non-binary, 1 other, 1 preferred not to say, and 1 missing) and 64 Oas (Mean_age_= 65.6 years, SD_age_= 5.15 year; 36 female, 28 male).

#### Materials

To accommodate the demands of the online testing environment, we reduced the number of trials per person to 20 (10 semantic trials and 10 phonological trials). This allowed the session to last approximately 30 minutes to ensure maximal participant engagement given the remote platform. A subset of 20 lists was selected for the current experiment from the initial pool of 46 theme lists used in Experiments 1 and 2 (see **Appendix D**). From these, we generated 12 counterbalanced versions of the experiment. Across versions, each list was assigned to the semantic condition in six versions and to the phonological condition in the remaining six versions, ensuring that each list appeared equally often in each condition across participants. In addition, the presentation order of the three themes within each list was rotated across versions using all six possible permutations, with each permutation in each sublist position occurring twice across the 12 versions. This ensured that neither condition assignment nor theme order was confounded with study list.

Recognition memory was assessed using a 12-item test following each study list. The test consisted of nine studied words (old items) and the three non-presented theme words (new items). To select these new test items, one of the four studied associates from each sublist was omitted from the test, leaving three studied associates to serve as old items. The omitted study position was counterbalanced across lists such that each serial position was omitted an equal number of times across the stimulus set. Finally, the nine old items and three new items were randomly intermixed for presentation during the recognition test.

#### Procedures

The experiment was programmed in PsychoPy (Version 2024.2.1) and administered online using Pavlovia (https://pavlovia.org/). Participants provided informed consent electronically in accordance with the University of California, Riverside Institutional Review Board (IRB) prior to beginning the study. To help with bot detection, before the experiment began, participants completed a text-based CAPTCHA in which they entered a five-character alphanumeric code displayed on the screen. Participants were given up to three attempts to enter the code correctly; failure on all three attempts resulted in termination of the experiment.

Following successful completion of the CAPTCHA, participants received detailed instructions about the task and how to respond. Participants were provided with an example of the task and then completed a comprehension check assessing their understanding of the task and the response mapping. Specifically, participants were asked what they should do if they believed they had previously seen an item during the study phase and were required to correctly indicate that they should select the button corresponding to the “old” response.

The study phase was identical to the unimodal presentation in Experiment 2: For each 12- item study list, words were presented visually only, one at a time, in the center of the screen. Each word remained on the screen for 1000 ms, with a 32 ms interstimulus interval between successive words. Following presentation of the study list, a screen indicating that the test was about to begin appeared on the screen for 3000 ms. Following this, the screen advanced automatically to the test phase in which the 12 test words were presented serially, one at a time, in the center of the screen. For each word, participants indicated whether the item had appeared in the preceding study list (“old”) or had not appeared (“new”). Responses were made by pressing the left or right arrow key. Participants had up to 3000 ms to make each response (timing based on Flegal et al., 2010) and labels remained on the screen until a response was made or the trial timed out to reduce non-task-related memory demands. To control for potential response biases, the assignment of the left and right arrow keys to the “old” and “new” response options was randomized across participants, with the mapping determined at the start of the experiment and remaining constant throughout the task. Immediately following the test phase and before beginning the next study phase, participants completed a simple arithmetic equation as an attention check. Participants were excluded from analysis if they scored below 75% on the math check or failed to provide responses on more than 50% of trials.

#### Data analysis

To assess recognition of studied items, analyses were restricted to trials for which the correct response was “old”. Hit rate was calculated as the number of correctly identified studied items divided by the total number of studied-item trials (hit rate = number of old items correctly identified as “old” / number of studied items), separately for each list condition (phonological, semantic) and sublist position (A, B, C); no-response trials were excluded from all analyses and calculations. A mixed-design repeated-measures ANOVA was conducted with age group (YAs, OAs) as a between-subjects factor and condition (phonological, semantic) and sublist (A, B, C) as within-subjects factors. Significant main effects were followed up using estimated marginal means and pairwise comparisons; Holm corrections were applied to the pairwise comparisons.

To examine whether memory for studied items varied as a function of study–test lag, analyses were restricted to studied-item (i.e., old) trials. Recognition responses were coded at the trial level as correct or incorrect, such that the model estimated the probability of a hit (i.e., correct identification of a studied item). Study–test lag was calculated for each studied item as the number of intervening items between its presentation during encoding and its corresponding presentation at test. Study–test lag was treated as a continuous variable, rather than being binned into categories, allowing us to test whether the probability of correctly recognizing a studied item changed systematically as a function of increasing lag. We used a binomial generalized linear mixed-effects model rather than an ANOVA, because hits are binary at the trial-level. This analysis included age group, condition, and z-scored study–test lag, all two- and three-way interactions as fixed effects, and participant as a random intercept. Study–test lag was z-scored to place the continuous predictor on a standardized scale, which aided model estimation and allowed effects involving lag to be expressed in standard-deviation units.

To assess false alarms (i.e., false memories) to new items, analyses were restricted to trials for which the correct response was “new.” False-alarm rate was calculated as the number of new items incorrectly identified as “old” divided by the total number of new-item trials. False-alarm rates were calculated separately for each participant, list condition (phonological, semantic), and lure sublist (A, B, C). A mixed-design repeated-measures ANOVA was then conducted with age group (YAs, OAs) as a between-subjects factor and condition (phonological, semantic) and sublist (A, B, C) as within-subjects factors. Significant effects were followed up using estimated marginal means and pairwise comparisons; Holm corrections were applied to pairwise comparisons.

For completeness, we report raw descriptive statistics of all response measures in **Table 6**.

**Table 6.**
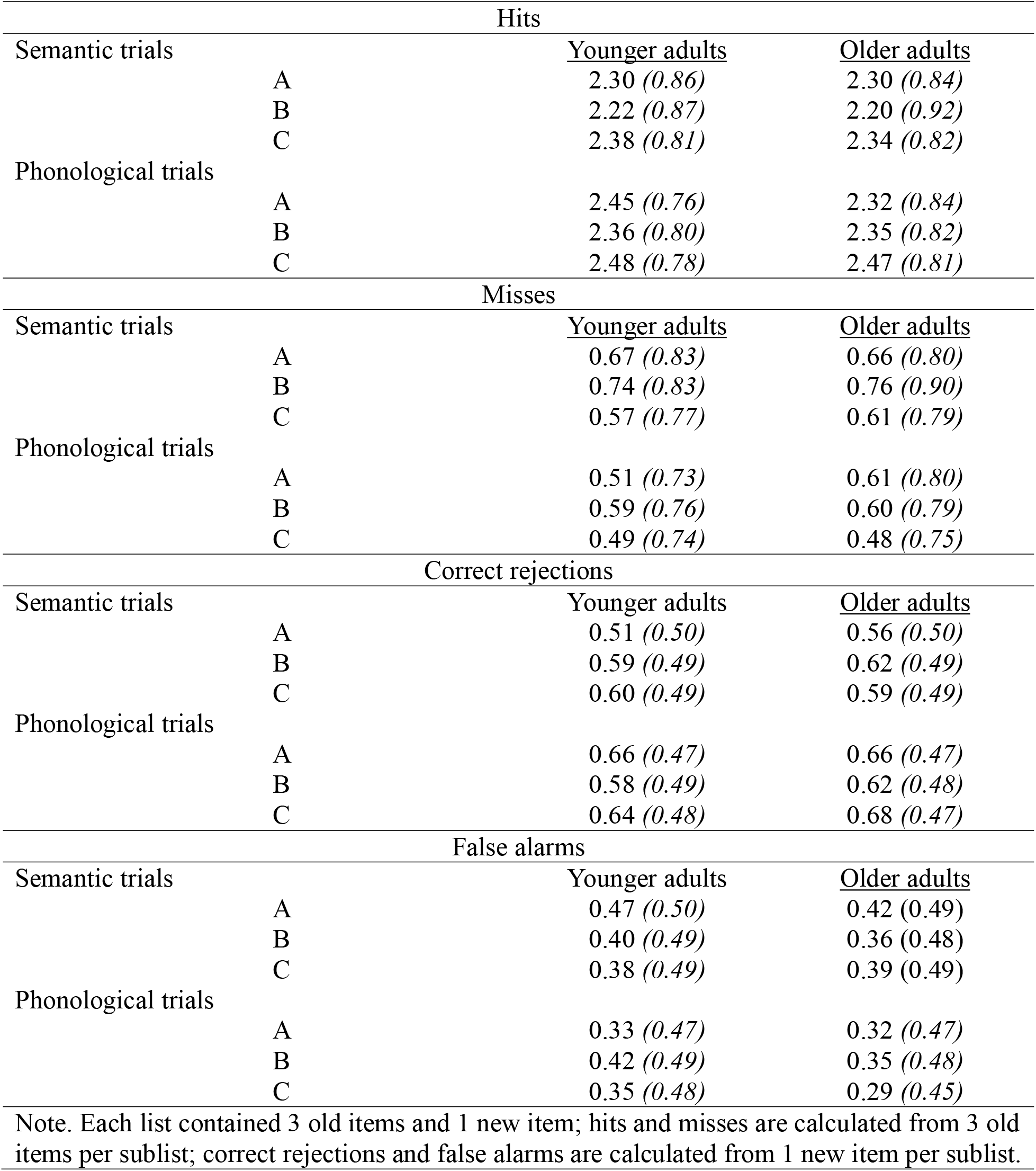
Average recognition responses (standard deviation in parentheses) split by age group, list type, and sublist for Experiment 3.

### Results

#### Correct recognition (hit rates)

Hit rates were analyzed using a 2 (age group: YAs, OAs) x 2 (condition: phonological, semantic) x 3 (sublist: A, B, C) mixed-design ANOVA, with age group as a between-subjects factor and condition and sublist as within-subjects factors. There was a significant main effect of condition (*F*(1, 126) = 31.91, *p* < .001, η²p = .202) with higher hit rates for phonological than semantic lists. There was also a significant main effect of sublist (*F*(1.97, 248.26) = 16.60, *p* < .001, η²p = .116). Follow-up comparisons indicated that hit rates were higher in sublist A compared to sublist B (*p* = .012) and higher in sublist C than in either sublist A (*p* = .006) or B (*p* < .001). The main effect of age group was not significant (*F*(1, 126) = 0.40, *p* = .530, η²p = .003), nor was the age group x condition interaction (*F*(1, 126) = 0.52, *p* = .472, η²p = .004), the age group x sublist interaction (*F*(1.97, 248.26) = 0.70, *p* = .497, η²p = .006), the condition x sublist interaction (*F*(1.98, 249.25) = 1.10, *p* = .334, η²p = .009), nor the 3-way interaction between age group, condition, and sublist (*F*(1.98, 249.25) = 2.63, *p* = .074, η²p = .020). Overall, these results indicate that participants showed higher hit rates for phonological than semantic lists and that hit rates varied across sublists; importantly, these effects did not differ significantly between YAs and OAs^2^. See **Table 7** for detailed descriptive statistics.

**Table 7.** Average hit and false alarm rates (standard deviation in parentheses) split by age group, list type, and sublist for Experiment 3.

| Hit rate |  |  |  |
| --- | --- | --- | --- |
| Semantic trials |  | <u>Younger adults</u> | <u>Older adults</u> |
|  | A | 0.77 (0.13) | 0.77 (0.12) |
|  | B | 0.74 (0.13) | 0.73 (0.14) |
|  | C | 0.79 (0.14) | 0.78 (0.14) |
| Phonological trials | A | 0.82 (0.13) | 0.77 (0.12) |
|  | B | 0.79 (0.12) | 0.78 (0.12) |
|  | C | 0.83 (0.14) | 0.82 (0.11) |
| False alarm rate |  |  |  |
| Semantic trials |  | <u>Younger adults</u> | <u>Older adults</u> |
|  | A | 0.47 (0.18) | 0.42 (0.20) |
|  | B | 0.40 (0.19) | 0.36 (0.19) |
|  | C | 0.38 (0.21) | 0.39 (0.22) |
| Phonological trials | A | 0.33 (0.21) | 0.32 (0.19) |
|  | B | 0.41 (0.17) | 0.35 (0.16) |
|  | C | 0.35 (0.20) | 0.29 (0.20) |
| Note. Hit rate = proportion of 3 old items correctly identified as “old”; false alarm rate = proportion of 1 new item incorrectly identified as “old”. |  |  |  |

#### Hits across study-test lag

Hit probability across study-test lag was analyzed using a binomial generalized linear mixed-effects model (GLMM), with age group, condition, and study–test lag as fixed effects and a random intercept for participant. This analysis revealed a significant main effect of condition, χ²(1) = 33.46, *p* < .001, with higher hit probability in the phonological condition than in the semantic condition (odds ratio = 1.27, *p* < .001). There was no significant effect of study–test lag (χ²(1) = 1.60, *p* = .205), indicating no evidence that hits declined as a function of lag. The age group x lag interaction (χ²(1) = 1.38, *p* = .240), condition x lag interaction (χ²(1) = 0.94, *p* = .331) and age group x condition x lag interaction (χ²(1) = 2.52, *p* = .113) were also nonsignificant. See **Figure 5** for an illustration of the data.

**Figure 5.**
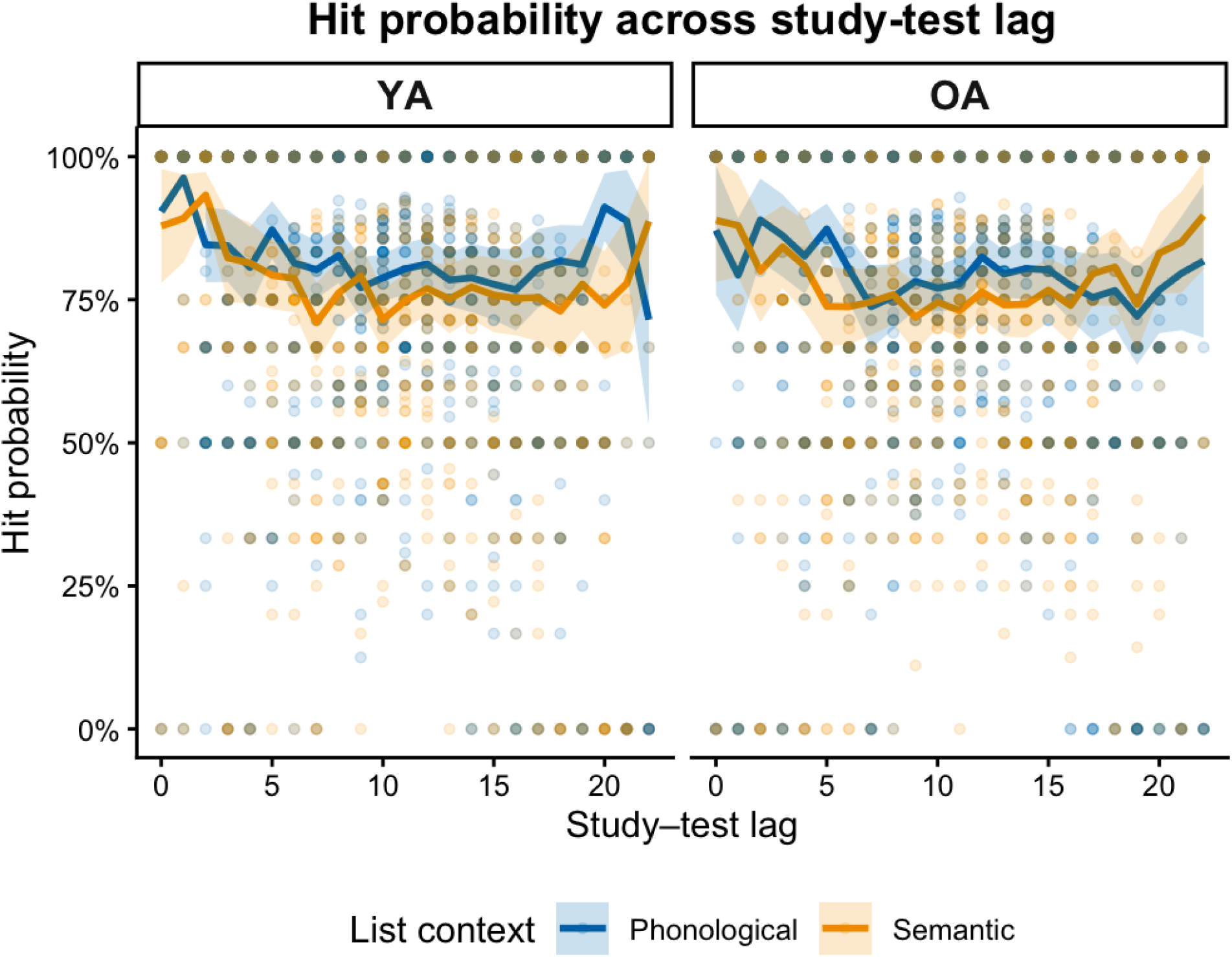
Hit probability as a function of study–test lag, age group (YA, OA), and condition (semantic, phonological). Hit probability was calculated as the proportion of correctly recognized studied items (“old” responses) at each lag for each participant. Points indicate individual participants, black points indicate group means, and error bars indicate 95% confidence intervals.

#### False memories (false alarm rate)

False alarm rates were analyzed using a 2 (age group: YAs, OAs) x 2 (condition: phonological, semantic) x 3 (sublist: A, B, C) mixed-design ANOVA. There was no significant main effect of age group (*F*(1, 126) = 0.85, *p* = .357, η²p = .007) and no significant age group x condition interaction (*F*(1, 126) = 0.01, *p* = .918, η²p = .001). There was a significant main effect of condition (*F*(1, 126) = 16.61, *p* < .001, η²p = .116) with lower false-alarm rates for phonological than semantic lists. There was also a significant main effect of sublist (*F*(1.97, 248.01) = 3.60, *p* = .029, η²p = .028). Follow-up comparisons indicated that false-alarm rates were significantly higher in sublist A than sublist C (*p* = .029), whereas the differences between sublist A and sublist B (*p* = .612) and between sublist B and sublist C (*p* = .114) were not significant. There was a significant condition x sublist interaction (*F*(1.91, 240.48) = 11.71, *p* < .001, η²p = .085). Follow-up analyses showed that false-alarm rates were significantly lower for phonological than semantic lists in sublist A (*p* < .001) and sublist C (*p* = .003) but did not differ between conditions in sublist B (*p* = .971). The age group x sublist interaction was not significant (*F*(1.97, 248.01) = 0.56, *p* = .569, η²p = .004) nor was the three-way age group x condition x sublist interaction (*F*(1.91, 240.48) = 1.90, *p* = .153, η²p = .015). Taken together, participants made fewer false alarms in the phonological compared to the semantic condition, particularly in sublists A and C, with no evidence that this pattern differed between YAs and OAs. See **Table 7** for detailed descriptive statistics.

## Experiment 3: Summary

In Experiment 3, we examined memory by providing additional retrieval support through an item-recognition test format. Participants showed better recognition performance for phonological than semantic lists, despite producing fewer correct recalls and more false memories for phonological than semantic lists in the two previous experiments. Additionally, hit rates were greater in sublist C than in either of the other sublists and greater in sublist A than in sublist B. Although we cannot compute serial-position curves from the item-recognition data, this pattern appears to parallel the strong recency effect and more moderate primacy effect observed in Experiments 1 and 2 (see Figure 2). Critically, these effects did not differ significantly between YAs and OAs. False-alarm rates were also lower for phonological than semantic lists, particularly for lures corresponding to the beginning and end of the lists (i.e., sublists A and C), with no evidence that this pattern differed as a function of age.

Taken together, findings from Experiments 1 and 2 do not support the prediction that the type of false memory should systematically vary with serial position as a function of differential reliance on gist and verbatim representations. Nevertheless, the recency advantage in veridical recall and the corresponding reduction in false memories for recently presented items remain partially compatible with FTT, insofar as greater availability of item-specific information may support accurate recall and reduce memory errors. The findings from Experiment 3 further suggest that providing additional retrieval support through recognition can alter how list composition influences memory performance, with these patterns remaining consistent across the lifespan. In particular, the advantage for phonological lists in recognition in Experiment 3, despite the disadvantage for phonological lists in free recall observed in Experiments 1 and 2, suggests that the effects of list composition may depend in part on the demands of the retrieval task. Recognition may provide sufficient retrieval support to allow participants to capitalize on the stronger representations associated with phonological lists, whereas free recall may place greater demands due to the self-generated nature of the retrieval task. Importantly, these patterns did not differ between YAs and OAs, suggesting that the influence of retrieval support and list composition on recognition performance is broadly preserved across the lifespan.

However, we cannot rule out the possibility that differences in testing format (i.e., in-person for Experiments 1 and 3 vs. online for Experiment 3), along with other concomitant differences between these individuals (Greene & Naveh-Benjamin, 2022), provide an alternative explanation for these effects of list composition. For example, one possibility may be that online participants engaged in shallower encoding which in turn could have influence how studied items were processed. Another possibility may be that online participants may have experienced greater environmental distraction or less standardized testing conditions, which could have influenced attention during the task. Thus, the observed differences between recall and recognition may reflect retrieval support, testing context, participant characteristics, or some combination of these factors and therefore findings should be interpreted cautiously.

## Discussion

This study examined the generation of false memories across putative short- and long-term memory stores, and how age-related changes influence these processes. Using a modified DRM paradigm with semantic and phonological sublists, in Experiment 1, we found that both younger adults (YAs) and older adults (OAs) were more likely to generate phonological than semantic false memories, regardless of sublist position. Although OAs recalled fewer items overall, the relative pattern of false memory generation was remarkably similar across age groups. In Experiment 2, which focused exclusively on OAs and restricted item presentation to the visual modality, we replicated key findings from Experiment 1. Once again, OAs generated a greater number of phonological as compared to semantic false memories, irrespective of list position. Critically, across both experiments, there was no interaction between sublist and list type, suggesting that semantic and phonological false memories can arise from all list positions. Experiment 3 extended these findings by examining recognition memory and revealed a related, but distinct, influence of list type: participants showed higher hit rates and lower false alarm rates in the phonologically- than semantically-related lists. Importantly, these effects did not differ as a function of age group. Intriguingly, changing the retrieval task from free recall to item recognition, thereby providing additional retrieval support seems to have altered the influence of list composition on memory. Whereas phonological lists produced more false memories during recall in Experiments 1 and 2, they produced fewer false memories and higher hit rates during recognition in Experiment 3.

Our findings from Experiments 1 and 2 align more closely with a shared-process account than with predictions derived from dual-store (Atkinson & Shiffrin, 1968; Glanzer & Cunitz, 1966) or Fuzzy Trace Theory (FTT; Brainerd & Reyna, 2002; Reyna & Brainerd, 1995) frameworks. Specifically, we did not observe the predicted dissociation whereby semantic false memories predominate in early list positions and phonological false memories in later positions, nor did OAs show a selective increase in semantic errors. Instead, semantic and phonological false memories were recalled across all list positions and age groups, suggesting that the nature of false memories is not strictly tied to the memory store from which they originate, but, instead, may reflect properties of the studied items in a list and list structure.

Notably, in Experiment 3, false alarm rates were lower for phonological than semantic lists in sublist A and sublist C. This pattern was unexpected given that phonological false memories were generally more prevalent than semantic false memories in the recall tasks in Experiment 1 and 2 and suggests that the influence of list position may interact with the type of similarity supporting false memory. Importantly, this pattern did not differ between YAs and OAs. Taken together, these findings further challenge the notion that reliance on semantic and phonological information systematically varies across list position. Instead, the variability in false memory effects across sublists is more consistent with a shared-process account in which the relative contribution of semantic and phonological similarity depends on the specific properties and organization of the studied lists. This interpretation is broadly consistent with related work showing that phonological similarity can strongly influence memory performance in working memory, whereas semantic similarity does not produce the same level of disruption (Kowialiewski et al., 2023). Further, the absence of age-related differences also suggests that the contribution of semantic and phonological similarity to false memory is more flexible and context-dependent than predicted by accounts proposing distinct memory stores.

### Shared cognitive processes underlying false memory generation

To better understand the mechanisms underlying these effects, it is important to consider the factors that may have contributed to the generation of false memories. FTT (Brainerd & Reyna, 2002; Reyna & Brainerd, 1995) suggests that information is simultaneously stored as both a verbatim and a gist representation which can both support accurate memory when these representations are strong. However, as verbatim traces decay more rapidly, individuals are thought to rely increasingly on gist representations, which, while more durable, are also more prone to errors, and, consequently, may contribute to the generation of false memories.

Consistent with this framework, both YAs and OAs in our study exhibited higher veridical recall for the most recent (i.e., sublist C) than earlier list positions. Notably, however, recognition of studied items in Experiment 3 did not decline as study–test lag increased. Although there appeared to be a numerically distinct pattern at the shorter study–test lag, with younger adults showing higher hit probability following phonological than semantic encoding and OAs showing the reverse pattern, this effect was not statistically significant. Thus, this suggests that the recall advantage for recently presented items may reflect greater accessibility during recall rather than a global loss of the memory representation for earlier items (for consistent evidence, see Cohen, 1970).

Additionally, during free recall, earlier items may also be subject to greater retrieval interference from subsequently presented items, with phonological similarity potentially introducing greater interference among related items. In contrast, during recognition, the studied items are shown explicitly at test, reducing the potential for interference. Moreover, because during recognition test items are presented visually, this format may have provided additional support for distinguishing phonologically related items based on their specific visual forms. This represents a source of information that is not available during free recall and may be particularly useful for distinguishing phonologically similar items for which surface-level visual information can help discriminate between otherwise similar-sounding words. Thus, the recognition advantage for phonological lists may reflect not only additional retrieval support but also differences in the information available at test.

FTT also predicts that semantic false memories should be more frequent for items presented at the beginning of a list, as these items are primarily supported by gist representations, which are more prone to distortions related to broader semantic content (Craik & Levy, 1970). In contrast, phonological false memories should be more frequent for items presented at the end of the list, where the verbatim representation, particularly of fragile surface-level information such as acoustic or phonological details, is strongest (Craik, 1968). Consistent with this, recent work (Coane et al., 2024) has shown that phonological false memories (measured as false alarms) are more frequent at short delays, whereas semantic false memories emerge more strongly after longer delays, suggesting that the contributions of verbatim and gist representations change over time and differentially influence false memory formation.

However, our findings did not support the above-mentioned prediction. Specifically, our data do not support the specific FTT prediction that the type of false memory should vary systematically with list position. However, the observed recency advantage in veridical recall is compatible with FTT’s distinction between verbatim and gist representations. One methodological difference that may account for this discrepancy with prior work from Coane and colleagues (2024) that used a recognition memory paradigm with relatively short lists (four or six items), probing primarily short-term memory processes, is that our study employed free recall and used substantially longer lists (twelve items), thereby engaging broader memory mechanisms. Additionally, in our recognition data, findings also did not show the predicted pattern of semantic false alarms being concentrated at earlier list positions and phonological false memories at later list positions. Instead, false-alarm rates showed a significant interaction between condition and lure sublist, with semantic false alarm rates higher than phonological false-alarm rates for lures corresponding to both the beginning (sublist A) and end (sublist C) of the list. Taken together, the absence of the predicted interaction between list type and sublist position in recall, together with the pattern of false alarms observed in recognition, suggests that differences in false memory type may not be solely attributable to differential reliance on putative short-term versus long-term memory stores.

One position-invariant factor that may contribute to false memory generation is source monitoring—i.e., the ability to accurately recall the context in which information was first learned (Johnson et al., 1993). Source monitoring is crucial for distinguishing between different memory sources and is largely supported by prefrontal regions (Johnson et al., 1993; Mitchell & Johnson, 2009). Studies with YAs show increased prefrontal activation in the short-term variant of the DRM task (Atkins & Reuter-Lorenz, 2011), highlighting the role of these regions in mitigating false memory generation. Although both age groups likely used source monitoring to adjudicate studied from falsely recalled items, OAs demonstrate well-documented deficits in source monitoring (Chalfonte & Johnson, 1996; Dehon & Brédart, 2004; Hashtroudi et al., 1989; Johnson et al., 1993; Mitchell et al., 2003; Mitchell & Johnson, 2009; Spencer & Raz, 1995) likely due to both structural and functional changes in frontal and core memory regions (Fjell et al., 2013; Salat et al., 2004).

To address potential concerns regarding age-related declines in executive control, OA participants in Experiment 1 and 2 were screened with a battery of neuropsychological measures (see **Table 1**), allowing us to at least partially mitigate concerns about deteriorations in executive control processing. Nevertheless, cognitive deficits in aging are multifactorial, encompassing impairments in memory discrimination (Korkki et al., 2020; Leal & Yassa, 2014; Stark et al., 2013), inhibitory control (Hasher et al., 1999), working memory deficits (Babcock & Salthouse, 1990; D’Esposito & Postle, 2015; Grady et al., 2006; Mattay et al., 2006), as well as greater difficulty in discriminating between externally and internally generated information (i.e., identifying whether a word was heard or thought about; Hashtroudi et al., 1989). Age-related deficits in source monitoring are particularly pronounced when the information being judged is highly similar (e.g., identifying which of two female speakers spoke; Ferguson et al., 1992). In our study, both the semantic and phonological conditions were designed to induce false memories; as such, source monitoring demands were, importantly, comparable across our conditions. Thus, the absence of age differences across false memory types in our study suggest that cognitive mechanisms supporting source monitoring operate similarly across semantic and phonological false memories, implying that shared cognitive processes contribute to both types of memory errors.

Aligning with this notion of shared cognitive mechanisms underlying both semantic and phonological false memories, the Activation-Monitoring Framework (AMF; Roediger et al., 2001) offers a parsimonious theoretical explanation for our findings (for a detailed discussion, see Chang & Brainerd, 2021). The AMF suggests that words may be connected by properties such as semantic or lexical networks as well as other properties such as acoustic or phonological features (Roediger et al., 2001). Accordingly, false memories arise through the automatic activation of related concepts, alongside source monitoring errors that lead individuals to misattribute the source of their memories^3^. This framework underscores how memory retrieval can be influenced by semantic associations and prior knowledge (Gallo & Roediger, 2003), as well as phonological similarity (Baddeley, 1964; Conrad, 1963; Wickelgren, 1965) and these semantic false memories may arise from spreading activation within semantic lexical networks (Collins & Loftus, 1975). Similarly, phonological similarity can lead to false memories by creating interference during recall, as demonstrated by Marmurek et al. (2006). Taken together, our findings suggest that false memories, whether semantic or phonological, emerge from shared processes, and highlight the importance of considering both phonological and semantic influences on memory retrieval within a unified theoretical framework for understanding memory distortions.

### Cognitive mechanisms underlying age differences in overall false memory rates

Although the relative pattern of false memory generation was similar across age groups, OAs exhibited higher overall rates of false memories in Experiment 1 and 2. Additional cognitive mechanisms may contribute to these age-related differences in overall false memory rates. Post-retrieval monitoring—evaluating the relevance of retrieved information to the goal of retrieval (Rugg, 2004)—declines with increasing age (Henkel et al., 1998; Johnson et al., 1993). Such monitoring processes are essential for verifying the veracity of retrieved information. In the context of the DRM paradigm, such monitoring failures may contribute to the increased susceptibility of older adults to false memories. In other words, OAs may be unable to monitor and reject a falsely remembered item that is similar to one they were originally presented with, thereby leaving them more prone to false memories in general (Gallo & Roediger, 2003; Mitchell & Johnson, 2009). Age-related declines in rehearsal and working memory may further reduce the ability of OAs to verify retrieved information and, thus, increase memory errors (for a further discussion of the importance of rehearsal in access of information, see Waugh & Norman, 1965).

Evidence indicates that OAs generate overall more false memories in working memory tasks relative to YAs (Jantz et al., 2021), possibly due to age-related declines in rehearsal processes (Naveh-Benjamin & Cowan, 2023). Relatedly, Pirolle and colleagues (2025) demonstrated that the rate of rehearsal is positively associated with performance, such that limiting rehearsal opportunities increases the incidence of false memories. Of note, despite the age-related differences observed in recall in Experiments 1, YAs and OAs showed similar patterns of recognition performance in Experiment 3, with no significant age differences in hit rates or false alarm rates. Taken together, these age-related declines in monitoring and rehearsal processes likely contribute to the elevated false memory rates we observed in OAs relative to YAs. That is, our findings suggest that while the type of false memory (semantic vs. phonological) may be governed by shared processes across age groups, the overall likelihood of false memory generation may be amplified in OAs during recall due to declines in monitoring and control processes.

Another factor that may contribute to group-level differences in the overall level of false memories in our recall data are the differences observed in veridical memory rates. In Experiment 1, we observed a significant main effect of age group on both correct responses and false memories. Although these age group effects did not interact with list type or sublist position, they raise the possibility that the differences observed in false memories could be explained by the differences in veridical memory performance. Additionally, this relative increase in the proportion of false memories in OAs was partly driven by lower recall of items overall in OAs (both veridical and false memories) compared to their younger counterparts. This aligns with previous findings that have reported overall reduced episodic memory even in cognitively healthy OAs (Brickman & Stern, 2009; Craik & Salthouse, 2011; Rönnlund et al., 2005). When focusing on correctly recalled items, OAs also recalled a lower proportion of correct items than YAs, consistent with prior research using the DRM paradigm in aging samples (Balota et al., 1999; Budson et al., 2003; Dennis et al., 2007; Norman & Schacter, 1997; Tun et al., 1998). Although we attempted to mitigate the impact of these group-level output differences by using proportion scoring (Khanna & Cortese, 2009; McGuire et al., 2015; Sugrue & Hayne, 2006), it could still be that differences in overall levels of false memories between age groups can at least be partially explained by differences in verbatim traces and concomitant veridical memory (c.f., Atkins & Reuter-Lorenz, 2008). Importantly, even when using proportion scoring to account for age-related differences in total output, there were no significant interactions between age and false memories with sublist position. Instead, we saw commonalities: Both YAs and OAs made more false memories in the phonological relative to the semantic condition and were more likely to make false memories from the A and B than C sublist.

These findings challenge the hypothesis that OAs would be disproportionately prone to semantic errors due to their presumed reliance on gist-based memory traces (Abadie & Camos, 2019; Brainerd et al., 2008; Grilli & Sheldon, 2022; Schacter et al., 1997). Importantly, the absence of a selective increase in semantic false memories in OAs challenges accounts that attribute age-related false memory increases primarily to enhanced reliance on gist-based representations. Instead, the comparable rates of phonological errors suggest that reduced access to item-specific or perceptual details suggest that false memory generation may be more influenced by factors such as the properties of the studied list, such as semantic or phonological similarity, rather than age-related memory differences.

Of note, Experiment 3 showed a different pattern: YAs and OAs did not differ in either hit or false memory rates. We speculate that this discrepancy between our recall and recognition findings across experiments may partly reflect differences in how memory was measured. Whereas Experiments 1 and 2 required participants to self-generate items during free recall, Experiment 3 required participants to make an old/new recognition judgment. Recognition and recall place different demands on memory retrieval and differences in measurement approaches can influence conclusions about memory performance (Brady et al., 2023). This interpretation is consistent with evidence that recall places greater demands on self-generated retrieval processes than recognition, which may contribute to larger age-related differences in recall (Craik & McDowd, 1987). Although older adults generally show poorer recognition than younger adults, the magnitude of this age difference is smaller than that observed for recall (Fraundorf et al., 2019; Rhodes et al., 2019), suggesting that differences between age groups may be more pronounced when memory must be retrieved without the support of the studied items. Thus, the absence of significant age group differences in the recognition measures examined here suggests that the age-related differences observed in recall in Experiments 1 and 2 may not generalize to recognition performance.

A further consideration is whether differences in participant characteristics contributed to the differing age effects across experiments. Research suggests online recruitment in cognitive-aging research may introduce selection effects, as OAs who participate in online research may differ from those typically recruited through community-based advertisements, particularly in characteristics related to technology access and literacy (Greene & Naveh-Benjamin, 2022). However, we did not collect neuropsychological measures with our online sample, limiting our ability to determine whether unmeasured participant characteristics contributed to differences across experiments. Thus, differences in recruitment and sample characteristics may represent an additional source of variability across experiments and should be considered when interpreting differences across the experiments.

### Role of list characteristics in false memory generation

Research beyond the aging literature also highlights the importance of list features in generating false memories (for an in-depth review, see Coane et al., 2021), highlighting roles of . associative strength (Kumar, 2021), categorical similarity (Buchanan et al., 1999), as well as orthographic or phonological similarity (Sommers & Lewis, 1999) in generation of false memories. Our findings extend this literature by demonstrating that list characteristics can fundamentally shape memory errors and the generation of false memories, irrespective of age. Critically, when items are presented in a way that encourages phonological errors, both YAs and OAs are equally susceptible to these types of memory distortions. This builds upon previous research by Dimsdale-Zucker (2018), who demonstrated that phonological errors can occur at all list positions. Importantly, the current study extends this work by showing that this effect occurs within the same participants, across age groups, and under the same study conditions.

Another key feature of the list design in our paradigm is the combination of a long, supra- span list—typically used to isolate memory representations from separate short- and long-term memory stores (Murdock, 1962)—with nested sublists that allow for more precise tagging of early, middle, and late study positions (adapted from Dimsdale-Zucker, et al., 2018). This structure allows us to examine the relative predominance of both veridical and false memories from different list positions. As expected, we observed a robust recency effect across both age groups in our recall data: YAs and OAs recalled a greater proportion of items from the end of the list (sublist C) than from earlier positions (sublists A and B). The recency advantage likely reflects the greater accessibility of recently encountered items, possibly due to reduced interference or decay (Glanzer & Cunitz, 1966; Murdock, 1962). Complementing this, the proportion of false memories was lowest for items presented at the end of the list (sublist C), in comparison to those from the beginning (sublist A) or middle (sublist B) positions. These results suggest that recent items are not only more accurately recalled but also less likely to result in false memories, possibly because they are stored as more accessible verbatim traces, minimizing effects of interference and decay. Experiment 3, however, suggests that the recency advantage may be more specific to free recall. Although recently presented items were more likely to be recalled, hit rates did not significantly vary as a function of study–test lag. This finding suggests that the influence of list position on memory may depend on the demands of the retrieval task. Overall, these findings indicate that the cognitive mechanisms underlying the recency effect remain largely intact across age, even in the presence of broader age-related declines in memory performance.

### Conclusion

From a theoretical perspective, our findings challenge the assumptions of traditional multi-store models (Atkinson & Shiffrin, 1968; Glanzer & Cunitz, 1966), which posit distinct memory systems, in explaining semantic and phonological false memories. In our first two experiments, we found that both YAs and OAs generated more phonological than semantic false memories, and, critically, that these errors occurred from across all serial positions. In our third, online pre-registered study using item recognition, false alarms occurred across all sublist positions, with lower false-alarm rates for phonological than semantic lists in sublists A and C. Our findings suggest that the cognitive mechanisms responsible for false memories might be more similar across short- and long-term memory stores than previously thought. That is, both semantic and phonological false memories may arise through common underlying cognitive processes, including the activation of related concepts and the misattribution of source information. This challenges the traditional dual-store perspective of memory (Atkinson & Shiffrin, 1968; Glanzer & Cunitz, 1966) and aligns more closely with the idea that memory is a fluid and interconnected system, where different types of information (e.g., phonological and semantic) are not processed or stored in isolation across different memory stores but rather coexist in a unified memory representation (Jonides et al., 2008; Nairne, 2002; however see Forsberg et al., 2025 for recent evidence suggesting an alternative perspective).

One consideration to note is that semantic and phonological lists consisted of different words, which could, in principle, introduce some variability in false memory rates across list types. Recent computational approaches (Petilli et al., 2025) offer methods to systematically manipulate semantic and phonological similarity within the same set of items, which could be applied in future work to further isolate the effects of similarity type and degree of similarity on false memory generation. Here, we counterbalanced which words were studied in the context of semantic versus phonological lists across participants to mitigate potential effects of list construction on our results.

Additionally, although we frame early versus late sublists as proxies for short-term versus long-term memory contributions, it is important to clarify that the present design does not allow for a strict separation of memory stores. Because each word was presented only once without a buffer task, items from early sublists could still have been actively maintained in working memory at recall (see also Waugh & Norman, 1965 for a discussion of transfer of information). Consistent with this interpretation, recognition performance did not decline systematically with increasing study–test lag, despite the robust recency effect observed in free recall, suggesting that temporal position does not map directly onto the availability of information in memory. Accordingly, sublist position effects are best interpreted as reflecting relative temporal position within the list rather than a pure short-term vs. long-term memory distinction.

In conclusion, this study provides important insights into the nature of false memory generation across the lifespan. While OAs exhibited lower veridical recall than YAs in Experiments 1 and 2, both age groups generated semantic and phonological false memories across list positions. In Experiment 3, YAs and OAs showed similar patterns of recognition performance and false-alarm rates with no evidence of age-related differences in the effects of condition or sublist. Together, our data suggest that age-related differences in memory performance may depend in part on retrieval demands, challenge the notion that semantic and phonological false memories operate via entirely separate storage mechanisms, and support the idea that partially overlapping cognitive processes drive these memory distortions.

## Declarations

### Ethics approval

This study was approved by the Boston College Ethics board.

### Consent to participate

Informed consent was obtained from all individual participants included in the study.

### Consent for publication

Informed consent was obtained from all individual participants included in the study for the publication of results derived from their participation.

### Availability of data and materials

The data and analysis scripts that support the findings of this study are available on https://github.com/dzmemoryandcontextlab/FMOA.git; only experiment 3 was preregistered (https://aspredicted.org/67k4i8.pdf; preregistration #304,917).

### Declaration of generative AI use

Claude AI (https://claude.ai/) was used to translate Python code into JavaScript compatible with the Pavlovia online testing environment for Experiment 3.

### CRediT

HDZ, EAK, and PAR contributed to Conceptualization. HDZ contributed to Investigation and Data curation. LG and HDZ contributed to Formal analysis. All authors contributed to Writing – original draft and Writing – review & editing.

## Acknowledgments

The authors thank the editor and reviewer for raising concerns that led us to conduct Experiment 3

## Appendix A. The forty-six possible study list triplets used in Experiments 1 & 2

|  |  |  |
| --- | --- | --- |
| buy-forever-nose | snake-blood-flash | sweet-doctor-pitch |
| bread-dirt-funny | whiskey-baby-handle | rough-morning-draw |
| church-man-urge | chain-vote-lesson | fog-basement-shake |
| finish-girl-process | beautiful-sleep-heaven | long-sneeze-train |
| site-boring-teeth | sheet-job-chair | rent-dance-fail |
| logic-sink-horse | high-math-dense | pen-annoy-hydrogen |
| shirt-broom-whole | hold-thief-boom | waiter-muscle-trash |
| window-give-impact | chaos-smart-height | slow-butterfly-justice |
| clam-fruit-data | smell-present-fence | king-bee-mine |
| focus-many-alone | back-roof-city | building-square-decrease |
| admission-ring-river | trip-basket-argue | marry-shovel-plain |
| soap-pig-nut | scale-fate-lie | army-dry-speaker |
| gas-piano-opinion | act-corn-hawk | atom-stress-hint |
| adult-anger-needle | black-cold-again | cheese-flag-lose |
| rain-mix-plaster | answer-letter-carpet | pan-bake-cabbage |
| rock-movie-saloon |  |  |

## Appendix B. The 138 theme words in alphabetical order and their phonological and semantic associates used in Experiments 1 & 2

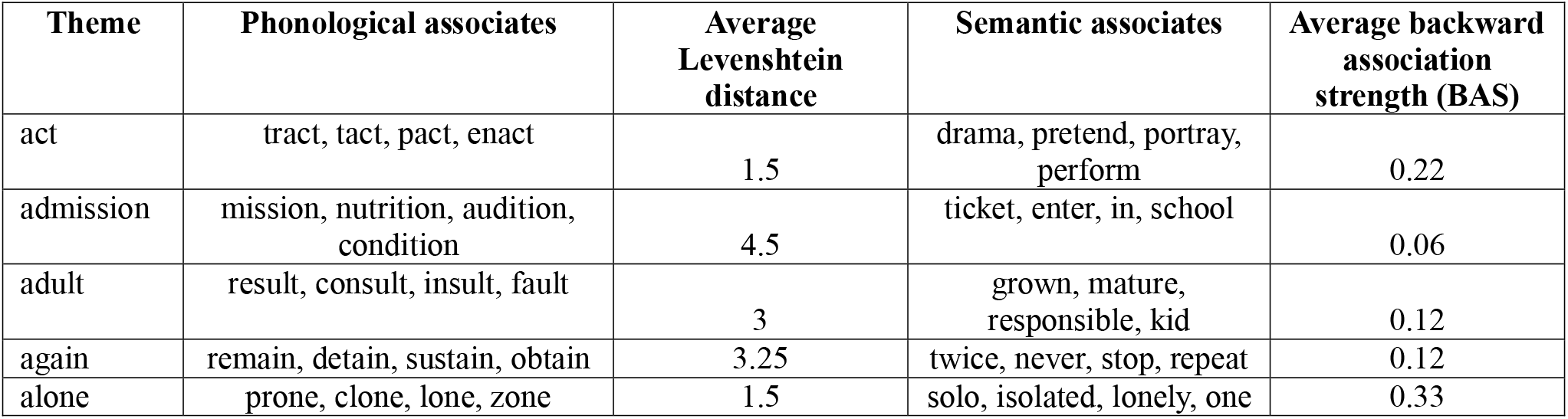
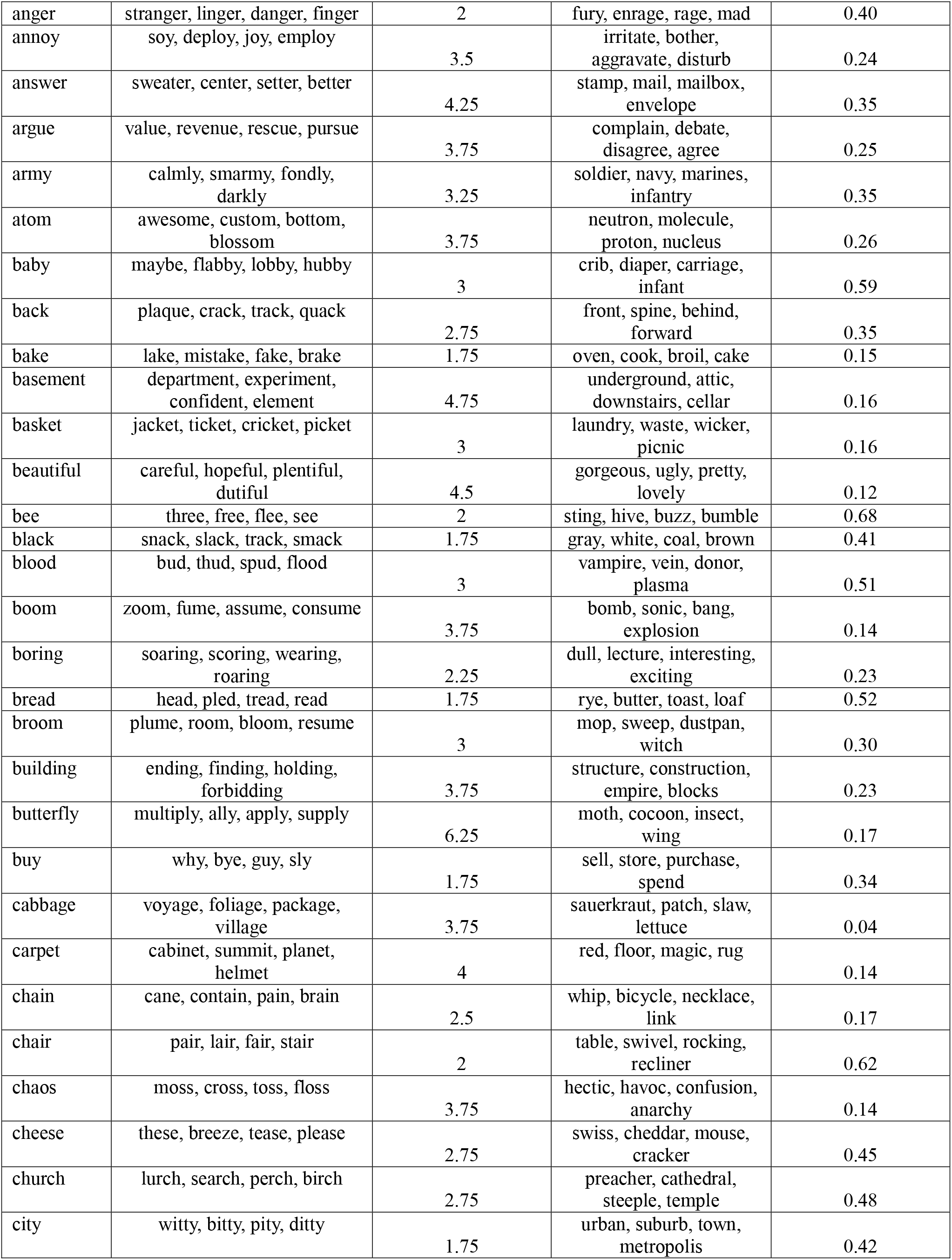
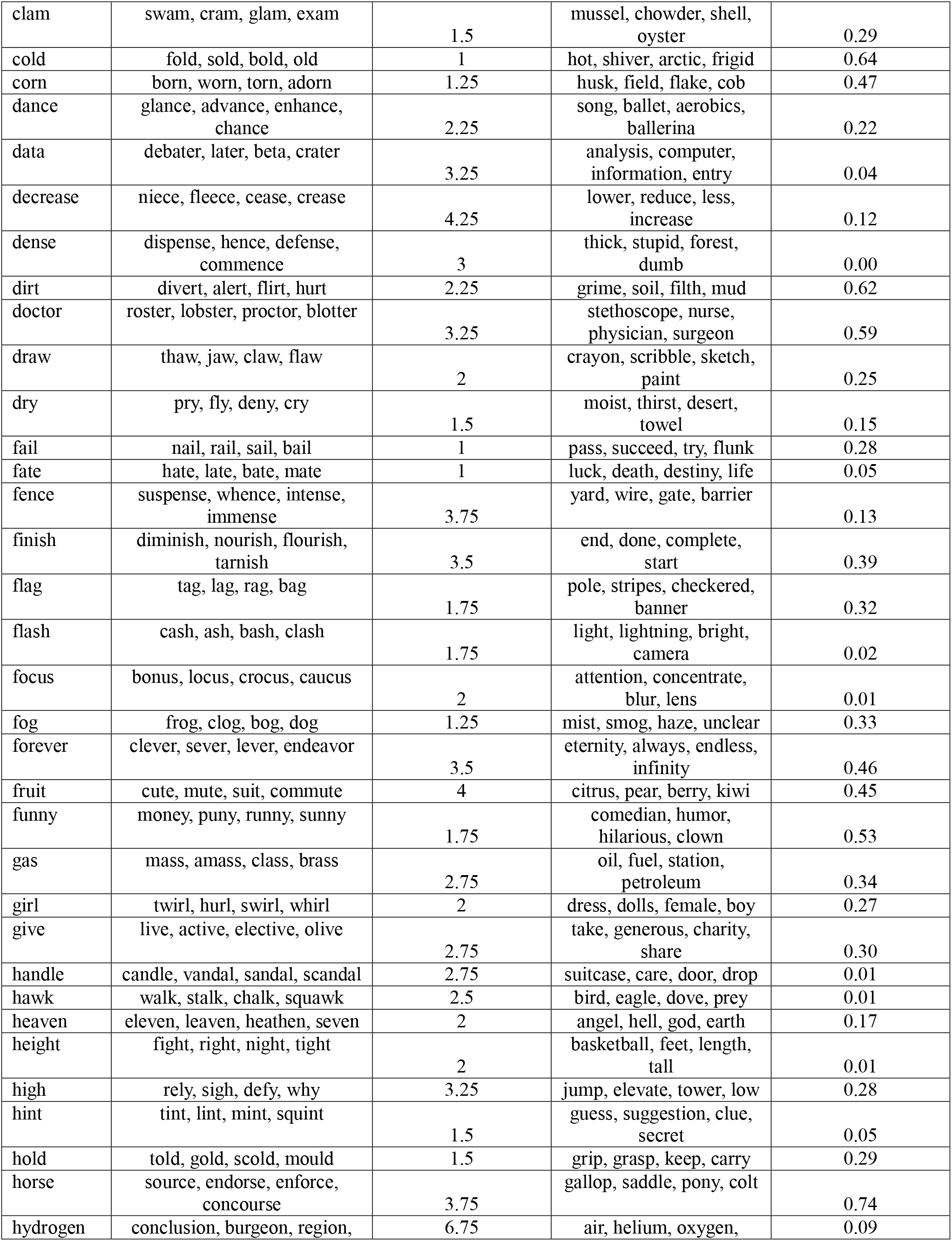
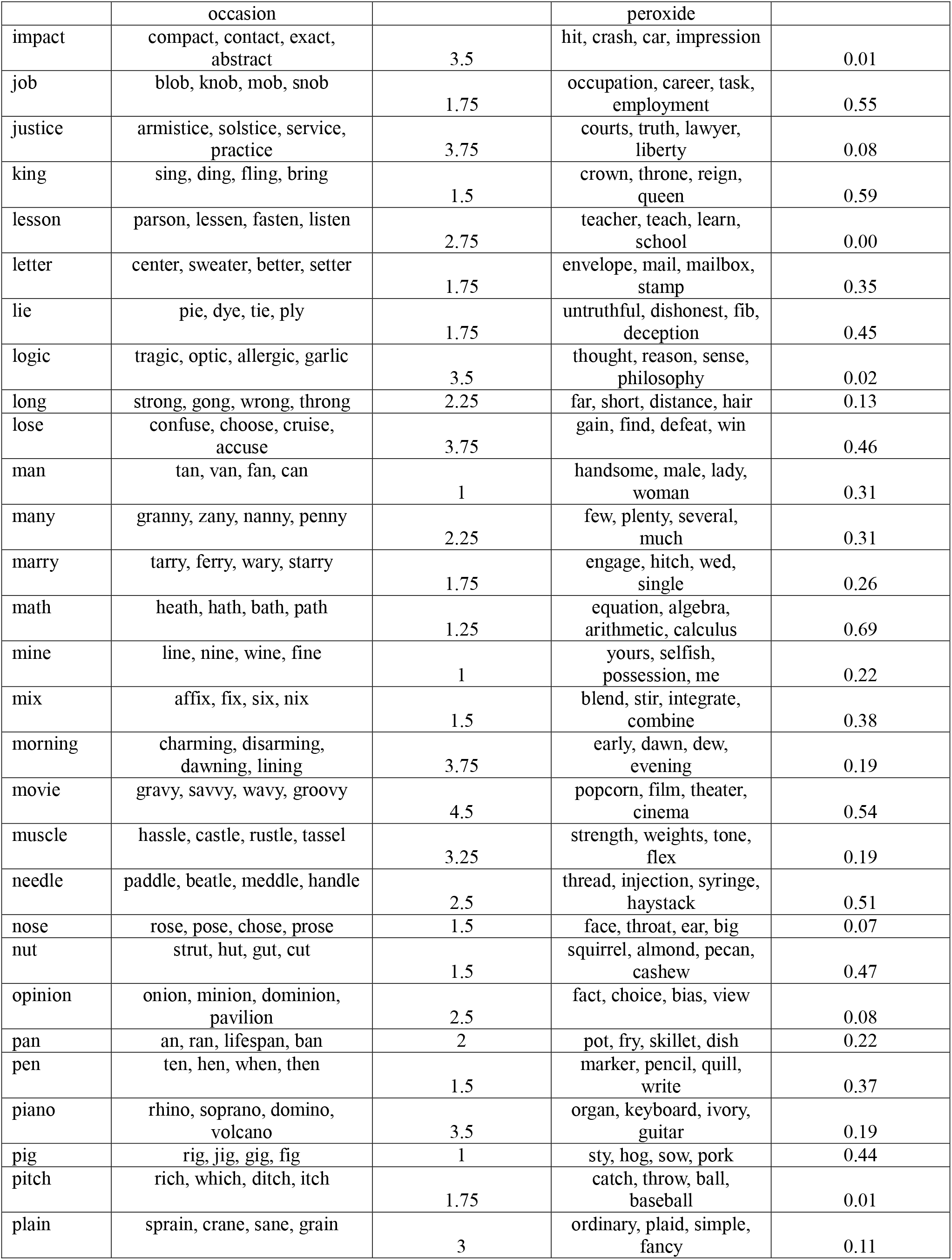
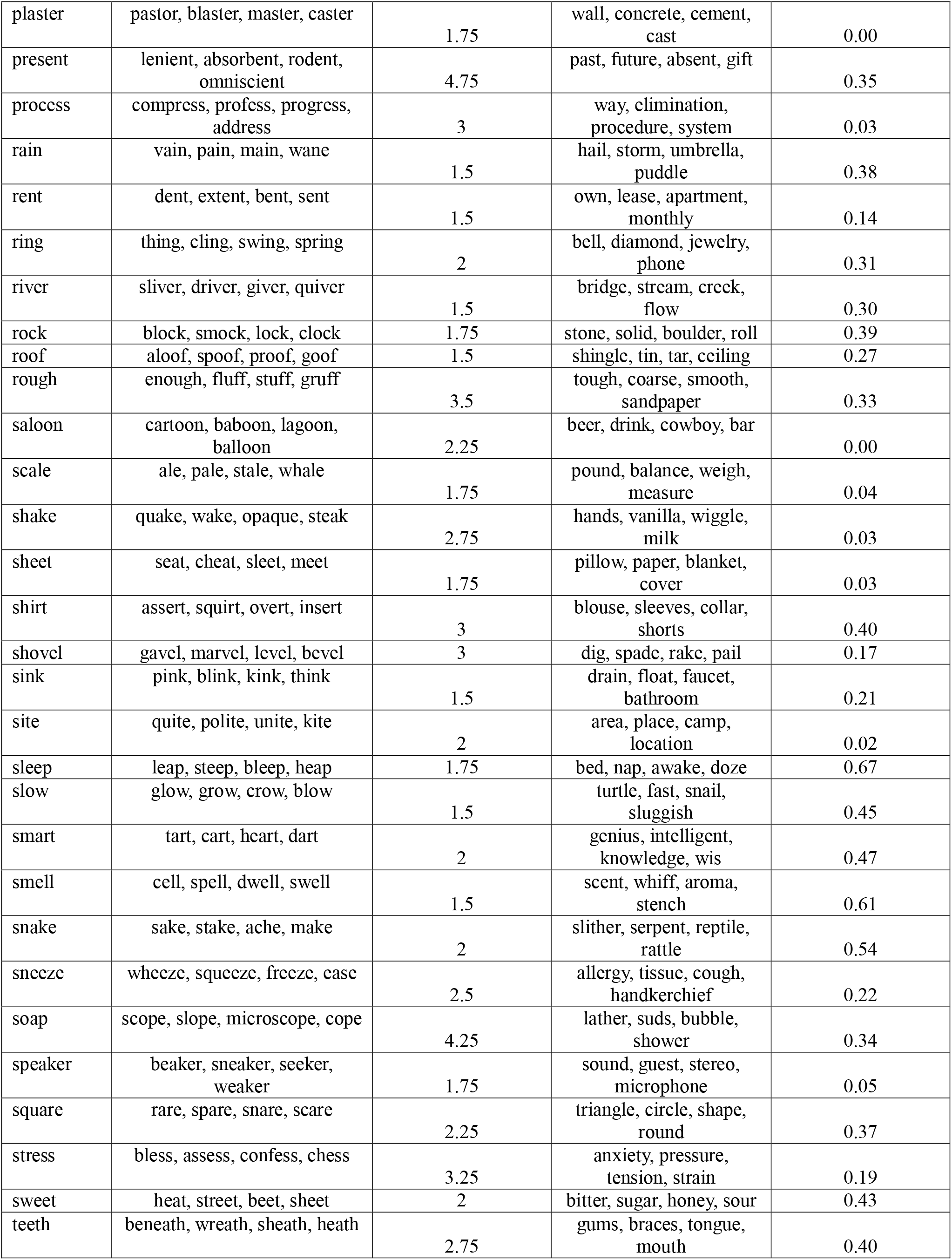
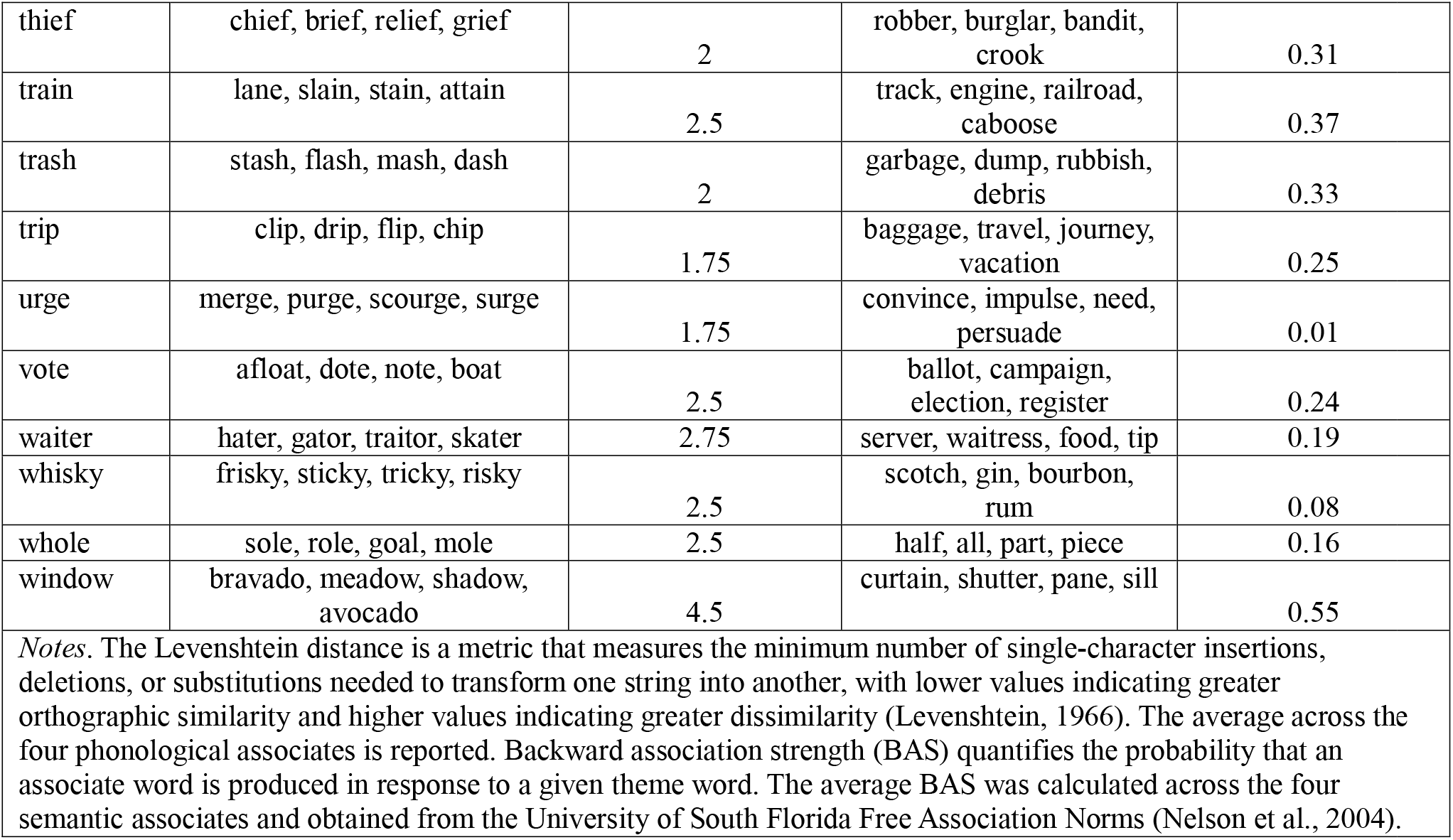

## Appendix C.

**Figure C1.**
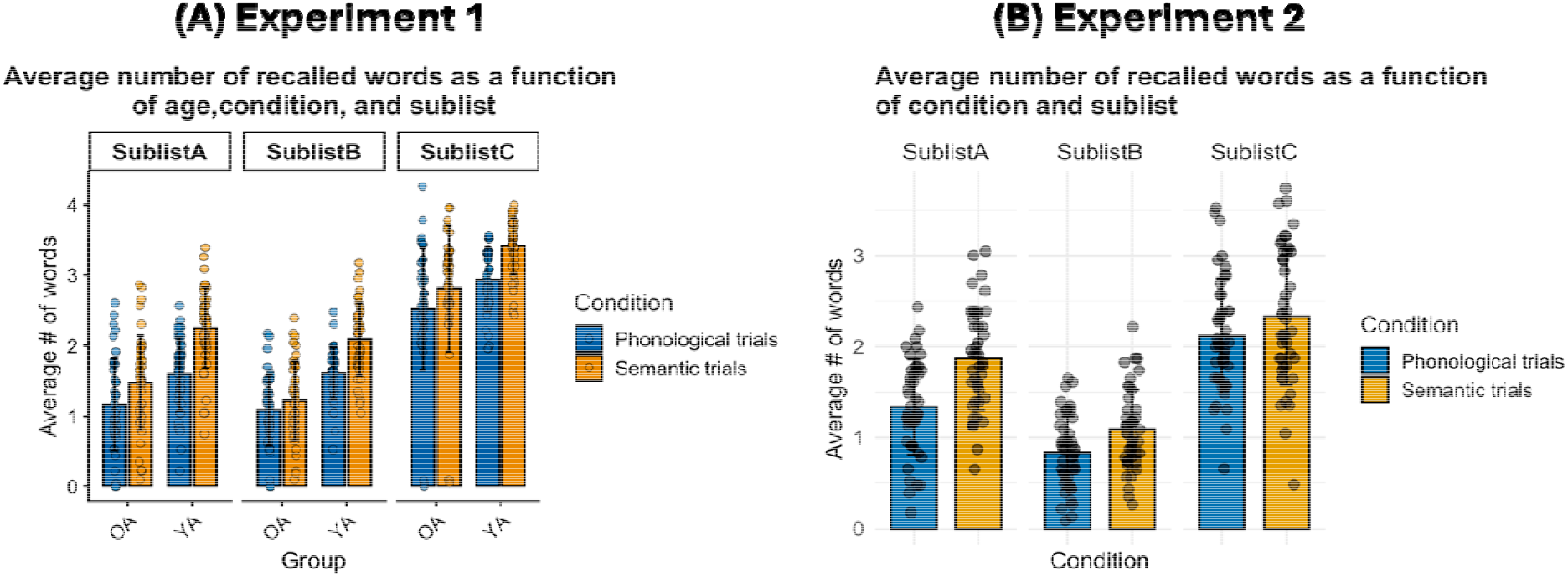
**(A)** Average number of words (correct and false) recalled by young adult (YA) and older adult (OA) participants as a function of condition (semantic, phonological) and sublist (A, B, C). **(B)** Average number of words recalled by OA participants as a function of condition (semantic, phonological) and sublist (A, B, C). Only older adults were tested in Experiment 2. Points indicate individual participants and error bars indicate ±1 standard deviation from the mean.

**Figure C2.**
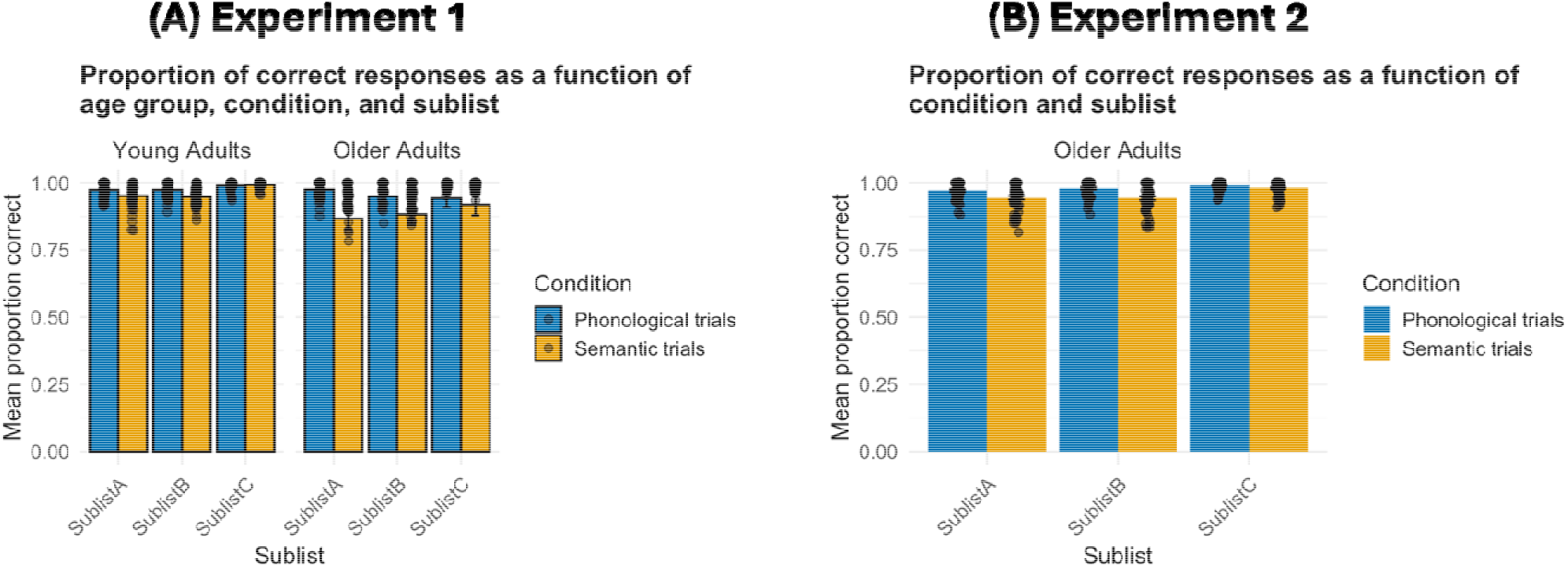
**(A)** Average proportions of correct responses recalled as a function of age group (YA, OA), condition (semantic, phonological), and sublist (A, B, C). **(B)** Proportion of correctly recalled words by older adult (OA) participants as a function of condition (semantic, phonological) and sublist (A, B, C). Only older adults were tested in Experiment 2. Points indicate individual participants and error bars indicate standard error of the mean.

## Appendix D. The 20 study list triplets used in Experiments 3

|  |  |
| --- | --- |
| window-give-impact | back-roof-city |
| admission-ring-river | trip-basket-argue |
| adult-anger-needle | scale-fate-lie |
| rain-mix-plaster | act-corn-hawk |
| whiskey-baby-handle | sweet-doctor-pitch |
| chain-vote-lesson | long-sneeze-train |
| beautiful-sleep-heaven | pen-annoy-hydrogen |
| sheet-job-chair | marry-shovel-plain |
| chaos-smart-height | atom-stress-hint |
| smell-present-fence | cheese-flag-lose |

## Notes

1 Although false memory in DRM paradigms is traditionally operationalized using only the critical lure, we also included other semantically related intrusions. This provides a more parsimonious and comprehensive index of false memory by capturing all opportunities for semantic false memories, following the approach used in Atkins & Reuter-Lorenz, 2008; Dimsdale-Zucker et al., 2019; MacDuffie et al., 2012.

2 Correct recognition rate (hits + correct rejections / total number of test trials) showed a similar pattern, with significant main effects of condition, *F*(1, 126) = 59.33, *p* < .001, η²p = .32, and sublist, *F*(1.98, 249.54) = 21.25, *p* < .001, η²p = .144. Here however, the age group x condition x sublist interaction was significant, *F*(1.98, 249.11) = 5.07, *p* = .007, η²p = .039. Follow-up analyses indicated that phonological lists had higher recognition accuracy than semantic lists across all age group x sublist combinations (Holm-adjusted *ps* ≤ .015). These results were consistent with the hit rate analysis in showing higher performance for phonological than semantic lists and variation across sublists. Although the age group x condition x sublist interaction was significant, follow-up comparisons showed that phonological lists yielded higher recognition accuracy than semantic lists across all sublist positions in both younger and older adults, indicating a broadly similar pattern across age groups.

3 Of note, these types of source monitoring errors are thought to stem from recollection-based processes—such as recall-to-reject and the distinctiveness heuristic (Gallo et al., 2006)–rather than from general executive control mechanisms as described by Johnson et al., (1993). However, the design of the present study was not intended to directly test the influence of these specific monitoring processes.

